# A speed-fidelity trade-off determines the mutation rate and virulence of an RNA virus

**DOI:** 10.1101/309880

**Authors:** William Fitzsimmons, Robert J. Woods, John T. McCrone, Andrew Woodman, Jamie J. Arnold, Madhumita Yennawar, Richard Evans, Craig E. Cameron, Adam S. Lauring

**Author notes:** Corresponding author Adam S. Lauring, 1150 W. Medical Center Dr. MSRB1 Room 5510B, Ann Arbor, MI 48109-5680.

## Abstract

Mutation rates can evolve through genetic drift, indirect selection due to genetic hitchhiking, or direct selection on the physicochemical cost of high fidelity. However, for many systems, it has been difficult to disentangle the relative impact of these forces empirically. In RNA viruses, an observed correlation between mutation rate and virulence has led many to argue that their extremely high mutation rates are advantageous, because they may allow for increased adaptability. This argument has profound implications, as it suggests that pathogenesis in many viral infections depends on rare or *de novo* mutations. Here we present data for an alternative model whereby RNA viruses evolve high mutation rates as a byproduct of selection for increased replicative speed. We find that a poliovirus antimutator, 3D^G64S^, has a significant replication defect and that wild type and 3D^G64S^ populations have similar adaptability in two distinct cellular environments. Experimental evolution of 3D^G64S^ under r-selection led to reversion and compensation of the fidelity phenotype. Mice infected with 3D^G64S^ exhibited delayed morbidity at doses well above the LD_50_, consistent with attenuation by slower growth as opposed to reduced mutational supply. Furthermore, compensation of the 3D^G64S^ growth defect restored virulence, while compensation of the fidelity phenotype did not. Our data are consistent with the kinetic proofreading model for biosynthetic reactions and suggest that speed is more important than accuracy. In contrast to what has been suggested for many RNA viruses, we find that within host spread is associated with viral replicative speed and not standing genetic diversity.

**Author Summary:** Mutation rate evolution has long been a fundamental problem in evolutionary biology. The polymerases of RNA viruses generally lack proofreading activity and exhibit extremely high mutation rates. Since most mutations are deleterious and mutation rates are tuned by natural selection, we asked why hasn’t the virus evolved to have a lower mutation rate? We used experimental evolution and a murine infection model to show that RNA virus mutation rates may actually be too high and are not necessarily adaptive. Rather, our data indicate that viral mutation rates are driven higher as a result of selection for viruses with faster replication kinetics. We suggest that viruses have high mutation rates, not because they facilitate adaption, but because it is hard to be both fast and accurate.

## Introduction

Mutation is the ultimate source of genetic variation, and mutation rates can have a significant impact on evolutionary rate [1–3]. The intraspecific variability in mutation rate in many viruses and bacteria indicates that mutation rates have been optimized by natural selection [4–13]. Given that most mutations are deleterious, the burden of excess mutational load will select against strains with abnormally high mutation rates [14–17]. This principle led to Sturtevant to ask, “Why does the mutation rate not evolve to zero?” [18,19].

A large body of theoretical and experimental work suggests that the selective pressure for higher mutation rates is due to either the physicochemical cost of maintaining a lower one or a selective advantage from an increased supply of beneficial mutations [20–23]. Many have argued for the adaptive benefit of high mutation rates in pathogenic microbes, which often exist in dynamic environments and are subject to host immune pressure [7,24,25]. However, direct selection of a variant with a higher mutation rate will only occur if it has been advantageous in the past, and in many cases, it has been difficult to separate the causes of a higher mutation rate from its consequences [19,26].

RNA viruses are ideal systems for studying the selective forces that act on mutation rates. While interspecific mutation rates range from 10^−4^ to 10^−6^ errors per nucleotide copied [4], studies of antimutators and hypermutators suggests that fidelity can only vary by several fold within a species [27]. The severe burden of mutational load exerts a strong downward pressure on mutation rates and hypermutator strains are attenuated *in vivo* [9,28–31]. Given the short generation times and remarkable fecundity of many RNA viruses, a small kinetic cost to higher fidelity should result in strong selection against antimutators [32,33]. However, the observed attenuation of antimutator RNA viruses *in vivo* has led many to argue for the adaptive benefit of high mutation rates, as genetic diversity provides a rich substrate for a virus’ evolution in the face of varying intrahost environments [7,10,34–38]. This concept is central to viral quasispecies theory, which generally proposes a link between genetic diversity and viral fitness [24,25].

Here, we define the selective forces that shape viral mutation rates by studying an antimutator variant. The 3D^G64S^ mutant of poliovirus was selected after serial passage in ribavirin, an RNA virus mutagen. The RNA dependent RNA polymerase (RdRp, 3D) of this variant contains a single glycine to serine substitution [5–7]. The basal mutation rate of 3D^G64S^ is reported to be ~20-30% that of wild type virus (WT). While the 3D^G64S^ mutant is attenuated in poliovirus receptor transgenic mice, the relative importance of replicative speed and fidelity to this phenotype is not clear [7,36]. Biochemical assays of 3D^G64S^ suggest a physicochemical cost of high fidelity, but as in other systems, its contribution to overall fitness remains unquantified [6,19,39].

## Results

We measured the relative fitness of 3D^G64S^ by direct competition over serial passage by RT-qPCR (Fig. 1A). Here, the fitness of 3D^G64S^ is 0.78 ± 0.01 (n=3 replicates) relative to WT. This is a moderate fitness defect, falling in the 64^th^ percentile in a dataset of 8970 fitness values obtained for point mutants of poliovirus under similar conditions [16] (e.g. HeLa cells, moi 0.1, 8 hour infection cycle, and 6 passages; Fig. 1B). We also measured the relative growth properties of WT and 3D^G64S^ using a plaque size assay, which measures the growth, burst size, and spread of individual viruses in the absence of competition [40–42]. The distribution of clonal plaque sizes was significantly different (p<0.005, unpaired t-test with Welch correction, n=272 WT and n=220 3D^G64S^ plaques), and consistent with a moderate fitness defect in 3D^G64S^ (Fig. 1C). In contrast to prior work, we were able to detect a significant replication defect for 3D^G64S^ by one step growth curve, but only with rigorous synchronization, more frequent time points, and larger numbers of replicates (Fig. 1D). This replication defect was not specific to HeLa, as we observed a similar lag for 3D^G64S^ in a 3T3 cell line that we derived from mouse embryonic fibroblasts from poliovirus receptor transgenic mice (PVR-3T3, Fig. 1E). These data demonstrate that the fitness defect of 3D^G64S^ is largely attributable to its slower replicative kinetics and is consistent with biochemical assays on purified RdRp [6,39].

**Figure 1:**
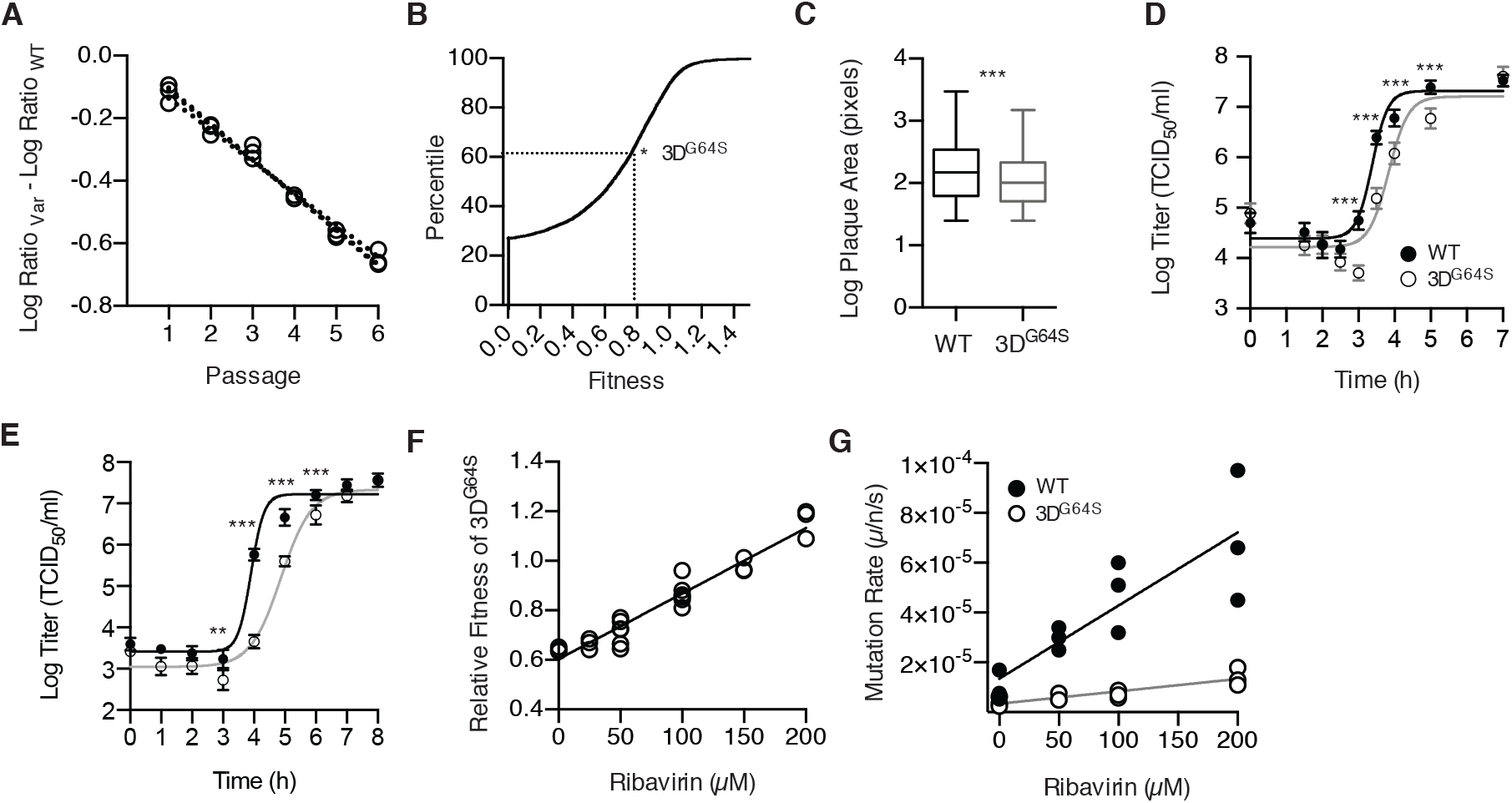
A speed-fidelity trade-off in the poliovirus RdRp. (A) Relative fitness of 3D^G64S^ as measured by direct competition. The amount of each virus at each passage was compared to the input and expressed as the difference in the log_10_ ratio in RNA genomes for 3D^G64S^ (open circles) relative to WT over time. The slope of the dotted lines are the relative fitness values, 0.78 ± 0.01, n=3 replicates. (B) Cumulative distribution function of fitness values for single nucleotide variants of poliovirus as determined in [16]. * indicates the relative fitness (0.78) and percentile (64^th^) of 3D^G64S^. (C) Plaque size of clones from WT (n=272, black) and 3D^G64S^ (n=220, grey) virus populations. Box plots show median, 25% and 75% quartiles, and 1.5x interquartile range. *** p ≤ 0.005, t-test with Welch’s correction. (D) Single cycle growth curve for WT (filled circles, black line) and 3D^G64S^ (open circles, grey line) in HeLa. Data are mean ± standard deviation (n=5 replicates). *** p<0.005, unpaired t-test comparing WT and 3D^G64S^ separately for each time point. (E) Single cycle growth curve for WT (filled circles, black line) and 3D^G64S^ (open circles, grey line) in 3T3 cell line derived from MEF of PVR transgenic mice. Data are mean ± standard deviation (n=5 replicates). ** p<0.01 *** p<0.005, unpaired t-test comparing WT and 3D^G64S^ separately for each time point. (F) Relative fitness of 3D^G64S^ (open circles) as measured by competition assay (see panel A) in the presence of varying concentrations of ribavirin. Note the baseline relative fitness of 3D^G64S^ (y-intercept) is lower than the fitness reported in panel A as the assays were performed under different experimental conditions (see methods). (G) Mutation rate in mutations per nucleotide per strand copied for WT (filled circles) and 3D^G64S^ (open circles) in the presence of varying concentrations of ribavirin, as determined by Luria Delbruck fluctuation test.

The reduced mutation rate and replicative fitness of 3D^G64S^ suggest a trade-off between speed and fidelity in RNA virus replication. Here, the fitness gain from increased replicative speed is offset by a reduction in fitness due to increased mutational load. We derived a quantitative model of this trade-off (see SI Model 1) by measuring the replicative fitness (Fig. 1F) and mutation rate (Fig. 1G, Table S1) of WT and 3D^G64S^ under exposure to an exogenous mutagen, ribavirin [43]. Wild type and 3D^G64S^ had equal fitness at approximately 150μM ribavirin. Based on these data, our model indicates that WT incurs a fitness cost of 0.137 from mutational load alone. Therefore, any fitness benefit of the high baseline mutation rates in WT would presumably need to offset this cost. In 3D^G64S^, the cost of mutational load is reduced to 0.037.

If viral RdRp are constrained by a speed-fidelity trade-off, selection for increased replicative speed (r-selection) will increase mutation rate. We subjected the 3D^G64S-1nt^ point mutant (A6176G) to r-selection over serial passage by infecting cells at low multiplicity and harvesting progeny at 4.5 hours (mid-exponential phase of replication). The 3D^G64S^ point mutant reverted to WT within 15 passages in 5 independent lineages (Fig. 2A). We only observed partial reversion at passage 15 in a subset of 24-hour control lineages, in which virus populations underwent twice as many cellular infection cycles per passage and experienced reduced r-selection. We next asked whether r-selection would lead to genetic compensation of the fidelity phenotype in 3D^G64S-3nt^, which has all three positions in the codon mutated to minimize reversion. After 50 passages of r-selection, we identified fixed and polymorphic single nucleotide variants (SNV) by next generation sequencing of all r-selected and control (24 hour passage) populations of 3D^G64S^ and WT (Fig. 2B).

**Figure 2.**
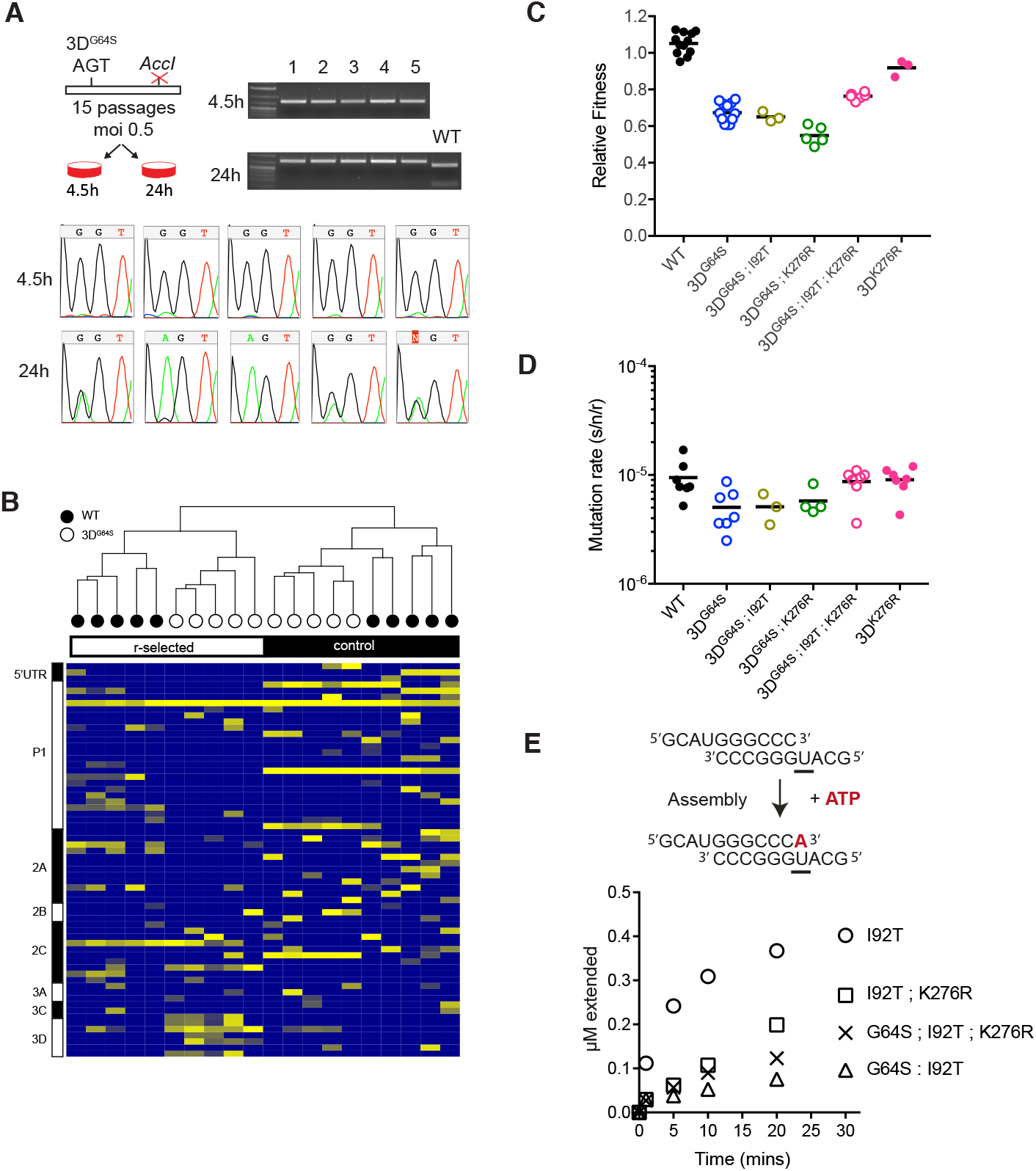
R-selection leads to increased mutation rates. (A) A point mutant of 3D^G64S^ (GGT^gly^ to AGT^ser^) was introduced into a poliovirus genome that is marked with a nearby point mutation that ablates an *AccI* restriction site. Viruses were serially passaged every 4.5 hours (r-selected) or every 24 hours (control) for 15 passages. Chromatograms show the codon for position 64 (either GGT^gly^ or AGT^ser^). Gel image of *AccI* restriction digest of all passage 15 populations showing that the reversion occurred in the parental backbone and was not due to contamination with WT virus, which retains the *AccI* site. (B) WT and a “locked in” version of 3D^G64S^ (GGT^gly^ to UCA^ser^) were subjected to r-selection (3.5-4 hour and 4-4.5 hour, respectively) or control (24 hour) passages for 50 passages as described in the text. Heatmap shows all mutations identified at >0.025 frequency in ≥2 out of the 20 total lineages, colored by log frequency. Diagram at left shows regions of the poliovirus genome. (C) Fitness of indicated variants relative to WT as determined by competition assay. Each symbol is a replicate competition assay and exact p-values for the key comparisons are provided in the main text. (D) Mutation rate of indicated variants in mutations per nucleotide per strand copied as determined by Luria Delbruck fluctuation test. Each symbol is a replicate fluctuation test and exact p-values for the key comparisons are provided in the main text. (E) In vitro kinetics of purified RdRp. Purified RdRp (2μM), primer-template (1μM) and ATP were incubated, and samples were quenched at the indicated time points (schematic). The kinetics of complex assembly and single-base incorporation are expressed as μM extended template (y-axis) over time (x-axis).

Unbiased hierarchical clustering of SNV by type and frequency indicates that the viruses explored distinct mutational pathways in adapting to either r-selective or control passaging regimes. Within the r-selected group, WT and 3D^G64S^ lineages clustered together and we noted a larger number of SNV within the coding region for the RdRp across the five 3D^G64S^ populations. Not surprisingly, we found that a number of distinct SNV increased viral fitness when introduced into the ancestral WT backbone. For example, the WT-VP4^S22G^ had a fitness of 1.53, and its presence in all r-selected and control lineages suggests that it mediates adaptation to HeLa cells. In contrast, a mutation in the viral helicase found only in r-selected populations, 2C^V127L^ (fitness 1.52-1.67 in WT and 1.11 ± 0.02 in 3D^G64S^), would be more likely to have a general effect on replicative speed.

To identify compensatory mutations, we focused our subsequent analysis on nonsynonymous mutations in the RdRp that were found predominantly in r-selected populations, shared among multiple lineages, and more frequent in 3D^G64S^ than in WT. Two mutations – U6261C/3D^I92T^ and A6813G/3D^K276R^ – met these criteria, and their frequencies at passages 30 and 50 suggest that the I92T mutation may have arisen first. The 3D^I92T^ mutation, which was found in both r-selected WT (3/5) and 3D^G64S^ (5/5) lineages, did not change either fitness or mutation rate appreciably in the 3D^G64S^ background (Fig. 2C and 2D). The r-selected K276R substitution, which was found in 3D^G64S^ lineages (4/5) and not in WT populations, decreased overall fitness in both WT (0.92 ± 0.03, p=0.0031 vs. WT, t-test) and 3D^G64S^ (0.55 ± 0.03, p=0.005 vs. 3D^G64S^, t-test). It had no detectable effect on mutation rate in either background. The G64S/I92T/K276R triple mutant had a significant increase in fitness (0.7637, p=0.0012 vs. 3D^G64S^, t-test) and mutation rate (8.71 x 10^−6^ s/n/r, p=0.0120 vs. 3D^G64S^, t-test) compared to 3D^G64S^ and each double mutant. Therefore, direct selection for replicative speed caused an increase in the poliovirus mutation rate with sign epistasis among the G64S, I92T, and K276R in the RdRp.

To gain mechanistic insight into the interactions among these three mutations, we analyzed the kinetics of single-base incorporation and misincorporation by purified RdRp. The 3D^G64S; I92T^ RdRp exhibits an assembly defect relative to 3D^I92T^ when incubated with purified primer-template and ATP (Fig. 2E and [6]). The K276R mutation partially compensates for this assembly defect in the 3D^G64S; I92T^ background, resulting in a 1.5-2 fold increase in incorporation of the correct nucleotide (A opposite U). This interaction is dependent on G64S, as K276R reduced RdRp activity in the 3D^I92T^ background. While some poliovirus mutators exhibit altered kinetics of base misincorporation for G opposite U [30], the kinetics of 3D^G64S; I92T; K276R^ were similar to those of 3D^G64S; I92T^ (Fig. S1).

The adaptability of WT and high fidelity viruses have generally been compared using assays that measure the acquisition of drug resistance, the reversion of an attenuating point mutation, or escape from microRNA in a limited number of replication cycles [5–7,34,36]. In these experiments, mutations come at little cost, and the assays essentially quantify the beneficial mutation rate. To capture better the impact of both deleterious and beneficial mutations on adaptability, we measured the fitness gain of WT and 3D^G64S^ over twenty passages in HeLa. While our WT strain is “culture-adapted,” we found that it was far from a fitness peak; both WT and 3D^G64S^ increased their fitness ten-fold in approximately 40 cellular infection cycles (20 passages, Fig. 3A, Fig. S2). The difference in the rate of fitness gain between WT and 3D^G64S^ lineages was small but statistically significant (0.025 per passage, WT > 3D^G64S^; mixed linear effects model, p=0.0129).

**Figure 3:**
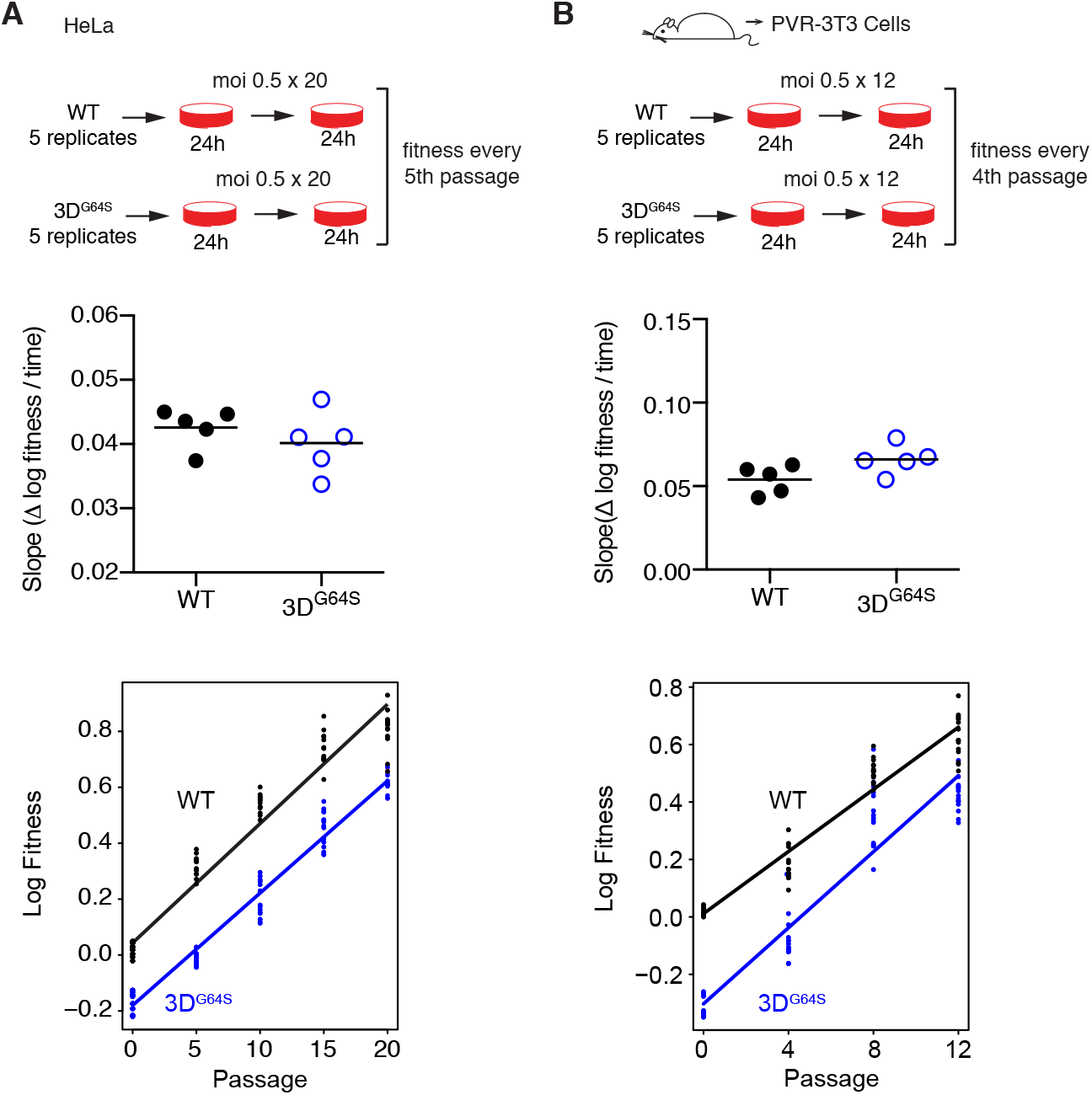
Adaptability of WT and 3D^G64S^ over 20 passages in HeLa (A) or 12 passages in a 3T3 cell line derived from mice transgenic for the poliovirus receptor (B). Fitness values (≥ 3 replicate competition assays for each data point) were determined for populations from every 5^th^ passage (A) or every 4^th^ passage (B) and the adaptability in the top panels expressed as the slope of the regression for log fitness over time for each of 5 independent lineages of WT (filled circles) and 3D^G64S^ (open circles, blue) for each cell line. The bottom panels show all the data from the 5 lineages together with the regression of log fitness over time. Exact p-values for the difference between the slopes for WT and 3D^G64S^ on HeLa (0.0129) and PVR-3T3 (0.0013) were derived from a mixed linear effects model (see Methods).

We examined adaptation to a completely distinct environment by repeating the experiment on our PVR-3T3 cell line [44,45]. In this alternative species and cell type, we actually observed greater fitness gain in the high fidelity 3D^G64S^ variant relative to WT (0.121 per passage; mixed linear effects model, p=0.0013). The larger fitness gain in 3T3 cells may reflect a larger supply of beneficial, compensatory mutations given the lower baseline fitness of 3D^G64S^ in these cells. These data suggest that, despite its two-fold reduction in mutation rate, 3D^G64S^ is not mutation limited, and that any adaptive benefit of a higher mutation rate is countered by the fitness cost of increased mutational load (see Fig. 1F and 1G and associated model above).

We next compared the phenotype of WT and 3D^G64S^ viruses *in vivo*, where the ability to generate genetic diversity may allow a virus to escape host immune restriction and to replicate better in a range of environments. Importantly, the available data have suggested that the attenuation of 3D^G64S^ and other high fidelity variants in experimental models is attributable to differences in the genetic diversity of the infecting population [7]. We therefore used next generation sequencing to compare the genetic diversity of 5 replicate stocks each of WT and 3D^G64S^. Using an internally benchmarked analysis pipeline that dramatically reduces false positive variant calls (see [46] and Methods), we identified no variants at greater than 0.1% frequency. Therefore, we can exclude any significant differences in standing genetic diversity between our WT and 3D^G64S^ populations. Even at extremely high multiplicities of infection, variants present at <0.1% are unlikely to complement each other or to cooperate reproducibly in a cellular context or in vivo [47].

The absence of mutational diversity in our replicate WT and 3D^G64S^ stocks is important, as poliovirus populations are subject to stringent bottleneck events, which further restrict intrahost diversity [48–50]. Work with barcoded RNA viruses suggests that the serial bottlenecks between the infecting population and the terminal population colonizing the central nervous system (CNS) are quite stringent [51,52], and we used published data to quantify the aggregate bottleneck size encountered by poliovirus in transgenic mice [45,48,49]. Maximum likelihood optimization of a simple probabilistic model estimated an aggregate bottleneck size of 2.67 between the inoculum and the CNS (Fig. 4A, SI Model 2). Therefore, the population that causes eventual disease in these mice is derived from no more than 2-3 viruses in the infecting population. In the setting of tight bottlenecks, many mutations will increase in frequency due to genetic drift as opposed to positive selection.

**Figure 4:**
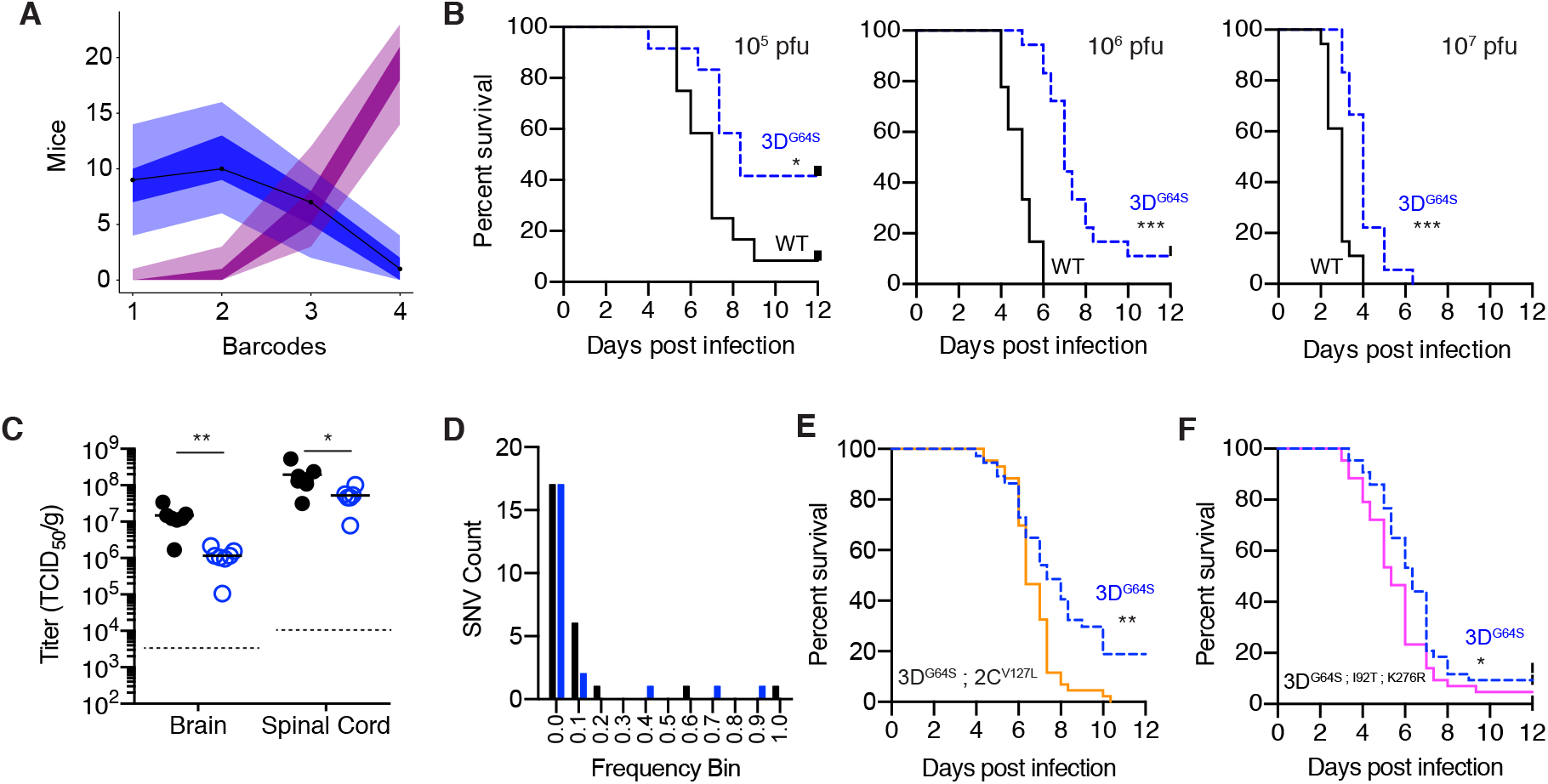
In vivo phenotype of WT and 3D^G64S^ (A) Maximum likelihood optimization of a simple binomial model (see Supporting Information Model 2) estimated an average inoculum to CNS bottleneck size of 2.67 (lambda 2.44, 95% CI 1.39-3.82) based on experimental data for 4 barcoded poliovirus populations [48]. Shown are outputs of 10,000 simulations of the model (number of mice with one, two, three, or four barcodes represented in the CNS). Each simulation represents 27 mice and each mouse has a bottleneck size drawn from a zero-truncated Poisson with an average lambda of 2.43 (blue) or 10 (magenta). Line is actual data from [48], the shaded regions represent the area occupied by 95% of the simulations, and the dark shaded regions representing the interquartile range of the simulations. (B) Survival curves showing mice with paralysis free survival over time for groups (n=18 per virus) infected intramuscularly with 10^5^ pfu (left), 10^6^ pfu (center), and 10^7^ pfu (right) of WT (black) or 3D^G64S^ (dashed blue). * p<0.05, *** p<0.001 by Log rank test. (C) Viral titer in brain and spinal cord 5 days post intravenous inoculation with 10^7^ pfu of WT (filled circles) or 3D^G64S^ (open circles). * p<0.05, **p<0.005 by Mann Whitney U test, n=7 mice in each group (out of 8 that were infected, one mouse in each group had titers below the limit of detection, dotted line). One mouse in each group had titers below the limit of detection (dotted lines). (D) Histogram of frequencies of intrahost SNV identified in the spinal cords of 12 mice from panel D (7 infected with WT and 5 infected with 3D^G64S^). Black, synonymous or noncoding; blue, nonsynonymous. (E) Survival curves showing mice with paralysis free survival over time for groups (n=43 per virus combined from two experiments) infected intramuscularly with 10^6^ pfu of 3D^G64S^ (dashed blue), or 3D^G64S^; 2C^V127L^ (orange). ** p<0.005, by Log rank test, actual p value 0.0012. (F) Survival curves showing mice with paralysis free survival over time for groups (n=43 per virus combined from two experiments) infected intramuscularly with 10^6^ pfu of 3D^G64S^ (dashed blue), or 3D^G64S; I92T; K276R^ (pink) *p<0.05 by Log rank test, actual p value 0.0411.

We infected groups of PVR mice intramuscularly with both WT and 3D^G64S^ populations. Both viruses were able to access the CNS efficiently through this route over a range of doses (Fig. 4B, Table S2), but there was a clear delay in the 3D^G64S^ group (p=0.0239, p<0.001, p<0.001 for 10^5^, 10^6^, 10^7^ pfu inocula, respectively). This lag persisted even when we increased the number of rare, but undetectable, variants in the inoculum by infecting with doses 20-fold higher than the LD_50_ of 3D^G64S^ (Table S3). Both viruses spread to the CNS and replicated to high titers after intravenous inoculation, although WT titers in the brain and spinal cord were marginally higher at 5 days post infection (Figure 4C, p=0.0012 for brain and p=0.0221 for spinal cord, Mann Whitney U test). Here too, WT mice generally developed severe morbidity more rapidly than those infected with 3D^G64S^. We characterized the mutations present in the CNS populations of 12 spinal cords of intravenously infected mice (Fig. 4D, 7 WT and 5 3D^G64S^). Most mutations were rare, none were shared among mice, and there was an excess of synonymous or noncoding variants relative to nonsynonymous ones. These data are consistent with random sampling of the infecting population as opposed to positive selection.

We also examined the impact of the r-selected mutations on virulence. The 2C^V127L^ mutation, which conferred a fitness of 1.11 in the 3D^G64S^ background and does not appear to affect fidelity (Fig. S3), restored virulence to nearly WT levels (p=0.0012, log rank test, compare 3D^G64S^ to 3D^G64S^;2C^V127L^ in Fig. 4E). In contrast, the triple mutant, 3D^G64S;I92T;K276R^, which replicates with a wild type mutation rate and a marginally increased fitness of 0.7637, was only slightly more virulent than the high fidelity variant 3D^G64S^(p=0.0411, log rank test, compare 3D^G64S^ to 3D^G64S;I92T;K276R^ in Fig. 4F). Therefore, restoration of replicative speed restored virulence in 3D^G64S^, but compensation of the fidelity phenotype did not.

## Discussion

We used a well-studied antimutator variant of poliovirus to identify the selective forces that optimize a pathogen’s mutation rate. Using three different assays, we identified a significant fitness cost to higher fidelity and directly link this cost to viral replication kinetics. Our quantitative model of the speed-fidelity trade-off suggests that selection for replicative speed has pushed viral mutation rates to a level that imposes a significant fitness cost at baseline due to lethal or highly deleterious mutations. Consistent with the trade-off model, direct selection for increased replicative speed led to indirect selection of polymerases with higher mutation rates. The genetic interactions are quite strong as the two compensatory mutations exhibited reciprocal sign epistasis. The speed-fidelity trade-off in the poliovirus RdRp appears to be a generalizable phenomenon, as compensatory mutations that increased the replicative speed of the 3D^K359H^ antimutator also increased its mutation rate (Table S4). Given the structural similarity among viral RdRp, the polymerases of other RNA viruses are likely to be subject to the same speed-fidelity trade-off, and we predict that the molecular mechanisms governing polymerase kinetics and mutation rate will be similar.

Trade-offs are essentially constraints that force one parameter to change with another. In this case, viral mutation rate changes with replicative speed [53]. Similar trade-offs are central to the kinetic proofreading hypothesis, which posits a close relationship between the error rates of biosynthetic processes and the kinetics of their component reactions [54]. Interestingly, studies of DNA replication and protein translation suggest that these systems optimize speed over accuracy, so long as the error rates are within a tolerable range [55,56]. We find a similar phenomenon in viral RdRp, where the WT generates an extraordinary amount of mutational load, largely because of the benefit in replicative speed.

Failure to consider evolutionary trade-offs can lead to teleological errors, in which the consequences of a process (e.g. increased genetic diversity) are misinterpreted as a cause (e.g. direct selection for a higher mutation rate [19,23,26]). Similarly, we find that the widely accepted link between within host genetic diversity and virulence is confounded by the fact that faster replicating viruses are both more virulent and have higher mutation rates. The high mutation rates of RNA viruses and the highly deleterious fitness effects of mutations ensure that most genetic diversity is extremely rare and unlikely to be consistently maintained in the face of intrahost and interhost bottlenecks [52]. We do not dispute that virus populations will harbor minority variants, that a subset of these mutations may be adaptive or beneficial to the virus, and that some may be virulence determinants. However, the observation of genetic diversity is not in and of itself evidence that selection has optimized mutation rates for the future benefit of novel mutations. Indeed, our data show little adaptive benefit to a marginally increased mutation rate and identify no plausible mechanism whereby the observed increase in rare genetic diversity can influence pathogenesis. We suspect that RNA viruses are subject to other trade-offs of evolutionary significance, perhaps between polymerase speed and recombination rate or recombination rate and polymerase fidelity. Here too, it will be important to define the selective forces at play, thereby separating the causes from the consequences.

## Methods

### Cells and viruses

A low passage stock of HeLa cells (<2 weeks in culture), previously obtained directly from ATCC (CCL-2), was kindly provided by Mary O’Riordan (University of Michigan). Except where noted, these cells were used for all experiments in this study and maintained in minimal essential media (MEM, Invitrogen 11090), supplemented with 10% fetal bovine serum (Gibco or Hyclone), 1x Penicillin-Streptomycin (Invitrogen 15140-148), 1x sodium pyruvate (Invitrogen 11360), 1x MEM alpha non-essential amino acids (Invitrogen 11140), 1x glutamine (Invitrogen 25030). A second stock of HeLa of unknown passage history was obtained from Michael Imperiale (University of Michigan). These cells were only used for plaque assays to titer stocks and were maintained in Dulbecco’s modified Eagle’s media (DMEM, Invitrogen 11965) supplemented with 10% fetal bovine serum and 1x Penicillin-Streptomycin. PVR-3T3 cells are described below and were maintained in DMEM supplemented with 10% fetal bovine serum, 1x Penicillin-Streptomycin and 1x glutamine. In all cases, cell lines were maintained for no more than 30 passages at a time. Wild type poliovirus and all mutants were generated from plasmid pEW-M, a Mahoney clone originally obtained from Eckard Wimmer (SUNY-Stonybrook) [57].

### Generation of PVR-3T3 cells

The University of Michigan Institutional Animal Care and Use Committee approved the protocols for the mouse studies described here and below. C57/BL6 PVR-Tg21 (PVR) mice [44,45] were obtained from S. Koike (Tokyo, Japan) via Julie Pfeiffer (UT Southwestern) and maintained in specific pathogen free conditions. Primary mouse embryonic fibroblasts (MEF) were derived from PVR mice. Day 13.5 embryos were harvested and washed in phosphate-buffered saline (PBS). The heads and viscera were removed, and the body was minced with a sterile razor blade, trypsinized, and homogenized by pipetting with a 10ml serological pipette. Cells were plated in DMEM supplemented with 10% fetal bovine serum, 1x Penicillin-Streptomycin and 1x glutamine. An immortalized cell was derived from PVR MEFs following the 3T3 protocol [58]. Briefly, freshly thawed MEFs were plated in 30 T25 flasks at a density of 3.8 x 10^5^ cells per flask in complete DMEM. Every third day, cells in each flask were trypsinized, counted, and transferred to fresh flasks at a density of 3.8 x 10^5^ cells per flask. As the cellular population began to increase (passages 13-15), cells were expanded into larger vessels and ultimately frozen down at passage 17.

### Site directed mutagenesis

All mutations were introduced into either pEW-M or subclones using overlap extension PCR [59]. The presence of the desired mutation and the absence of additional mutations were verified by Sanger sequencing of the amplified insert, and in some cases the entire genome.

### In vitro transcription, transfection, and viral stocks

Viral RNA was generated by in vitro transcription of the corresponding plasmid clone using T7 RNA polymerase, and virus was recovered following RNA transfection of HeLa. For transfections, 2.6 x 10^5^ HeLa were plated per well in a 12 well plate the day prior. One microgram of RNA was mixed with 4μl TransIT mRNA transfection reagent (Mirus 2225) and 100μl OptiMEM (Invitrogen 31985), incubated according to the manufacturer’s protocol, and applied to cells. Passage 0 virus was harvested at 100% CPE (within 24-48 hours). Passage 1 stocks were generated by passaging 100μl of passage 0 virus on fresh cells and were titered by either plaque assay or TCID_50_. Passage 2 and 3 stocks were generated by passaging at an MOI 0.01. For all stocks, cells were subjected to three freeze-thaw cycles and the supernatants clarified by centrifugation a 1400 x g for 4 minutes. These supernatants were stored at minus 80°C in aliquots to limit the number of subsequent freeze-thaw cycles.

### Competition assay for viral fitness

Competition assays were performed essentially as described in [17,42]. For the experiment in Fig. 1A, HeLa cells were plated in 12 well plates, at a density of 2.6 x 10^5^ per well the day prior to infection. Cells were infected at a total MOI of 0.1 with an equal TCID_50_ of WT and 3D^G64S^. Three replicate wells were infected with each pair of viruses in 250μl for one hour with occasional rocking. After one hour, the inoculum was removed and 1ml fresh media applied. Passage 1 virus was harvested after an additional 7 hours (8 hours since infection). The titer of the passage 1 virus was used to calculate the dilution factor necessary to maintain an MOI of 0.1 for the subsequent 5 passages. RNA was harvested from each passage using Trizol (Ambion 15596026). Random hexamers were used to prime cDNA synthesis with 1/10 of the RNA. Each cDNA was analyzed by real time PCR using three different primer and/or probe sets with duplicate PCR reactions for each sample/primer set. The first set, COM2F 5’ CATGGCAGCCCCGGAACAGG 3’ and COM2R 5’ TGTGATGGATCCGGGGGTAGCG 3’, was used to quantify total viral genomic RNA in a SYBR green reaction (Power SYBR Green PCR Master Mix, Thermo 4368708). Two custom TaqMan probes (Applied Biosystems) were used to quantify the number of WT and 3D^G64S^ genomes. Duplicate wells were averaged, and relative amounts of WT and 3D^G64S^ RNA were determined by normalizing the cycle thresholds for each these probes to those of the COM primer set (ΔCt = Ct_Virus_-Ct_GO_M). The normalized values for virus passages 1–6 were then compared to passage 0 to obtain a ratio relative to P0 (ΔΔCt = ΔCt_PX_-ΔCt_P0_). This relative Ct value was converted to reflect the fold change in the ratio (Δratio = 2^−ΔΔCt^). The change in ratio of the mutant relative to the change in ratio of the WT as a function of passage is the fitness ([Δlog ratio_Mut_-Δlog ratio_WT_]/time). Competition assays in ribavirin (Fig. 1F) were performed in the exact same manner except that serum free media were used in both drug and mock passages.

For all other competition assays (Fig. 3), we compared the experimental virus (e.g. WT P4, 3D^G64S^ P8 etc.) to a tagged WT reference (Tag8). We plated 2.6 x 10^5^ cells per well (either HeLa or PVR 3T3) in 12 well plates. Infections were performed at an MOI of 0.05 in 250μl complete media for 1 hour. After an hour, the media were aspirated and fresh 1ml growth media applied. All passages were for 24 hours. The dilution factor between passages required to maintain this MOI was 400 for HeLa competitions and 350 for PVR-3T3 competitions. All RNA harvests for these competitions were performed in 96 well plates using Purelink Pro 96 Viral RNA/DNA kits (Invitrogen 12280) and cDNA synthesis performed as above. In addition to the COM primer set (see above) we used primer pairs Tag8 seq.tag 5’ TTCAGCGTCAGGTTGTTGA 3’ + Rev. WT seq.tag 5’ CAGTGTTTGGGAGAGCGTCT 3’ and WT seq.tag 5’ AGCGTGCGCTTGTTGCGA 3’ + Rev. WT seq.tag 5’ CAGTGTTTGGGAGAGCGTCT 3’ to quantify the Tag8 reference and test samples respectively. Note also that in these competitions, the regressions were fit through passages 1–4 and excluded P0 as slight deviations from a 1:1 ratio of the two viruses in the inoculum can skew the slope when fit through this data point.

### Plaque size assay

Plaque assays were performed on subconfluent monolayers (7.5 x 10^6^ on day of infection) in 10cm dishes. The amount of virus applied to each plate was determined empirically to ensure well spaced plaques (~30 per 10cm dish). Plates were stained with crystal violet at 72 hours post infection. Each plate was scanned individually at 300 dpi using a flat-bed scanner. Sixteen bit image files were analyzed using ImageJ. Brightness, contrast, and circularity thresholds for plaque identification were set using uninfected plates.

### Single replication cycle growth curve

The day prior to infection, 4 x 10^5^ HeLa cells were plated in 12 well plates with 45 wells per virus (9 time points and 5 replicates per time point). Cells were infected at an MOI of 1 in 150μl volume and incubated on ice for 1 hour with occasional rocking. At one hour, the inocula were aspirated, each well was washed twice with ice cold PBS, and 1ml of fresh, pre-warmed growth media were applied to all wells. One set of 5 wells was immediately frozen as the t=0h time point. All other plates were returned to the incubator, and a set of 5 wells was removed and frozen at t=1.5, 2, 2.5, 3, 3.5, 4, 5, and 7 hours. All samples were titered by TCID_50_. The growth curve on PVR-3T3 cells was performed using a similar protocol, except that 5 x 10^5^ cells were plated the day prior and the time points were t=1, 2, 3, 4, 5, 6, 7, and 8 hours.

### Measurement of viral mutation rates

Mutation rates were measured by Luria-Delbruck fluctuation test, which in this case quantifies the rate at which the poliovirus 2C protein acquires the necessary point mutations to permit viral growth in 1mM guanidine hydrochloride [60–62]. Each fluctuation test was performed with 29 replicate cultures in 48 well plates. Sixty five thousand HeLa cells per well were plated the day prior to infection. In all cases, the media were changed to serum free media 3 hours prior to infection. For infections in ribavirin, this serum free media also included drug at the specified concentrations. Each well was infected in 200μl volume with 1000-4000 pfu per well depending on the virus and experimental condition. Five independent aliquots were also saved for subsequent titering (see N_i_ below). For infections in ribavirin, the infection media also included drug at the specified concentrations. Infected cells were incubated for 7 hours and then frozen. The lysed cells and media were harvested following three complete freeze-thaw cycles and transferred to a microcentrifuge tube. The empty wells were rinsed with 300μl of complete growth media and combined with the initial 200μl lysate. This 500μl lysate was clarified by centrifugation at 1400 x g for 4 minutes. Twenty-four wells were titered by plaque assay with 1mM guanidine hydrochloride in the overlay (see P_0_ below). Five wells were titered by standard plaque assay without guanidine hydrochloride (see N_f_ below). The mutation rate, μ_0_, was estimated from these data using the P_0_ null-class model: μ_0_ = −lnP_0_/(N_f_-N_i_), where P_0_ was the fraction of the cultures that yielded no guanidine resistant plaques, N_f_ was the average number of pfu in the absence of guanidine and Ni was the average number of pfu in the inoculum. As described in [63], μ_0_ can be converted to the mutation rate in nucleotide units by correcting for the mutation target (number of mutations leading to the scored phenotype, T) and the number of possible mutations at each target site (constant, 3) using the equation μ=3μ_0_/T. The number of distinct mutations that could yield the guanidine resistant phenotype was determined empirically by isolating and sequencing the entire 2C open reading frame for 15 guanidine resistant plaques derived from WT virus and 15 guanidine resistant plaques derived from WT virus treated with 200μM ribavirin. In each case we found 6 mutations that mediated resistance, although there were 7 total among 30 plaques (Table S1).

### Mutagen sensitivity assay

HeLa cells were plated the day prior to infection at a density of 2.6 x 10^5^ cells per well in a 12 well dish. On the day of infection, monolayers were pretreated with 0–600 μM ribavirin in serum-free media for 3 hours, then infected with virus at an MOI of 0.1 (50,000 pfu) for 60 min. The cells were washed once in phosphate buffered saline and incubated in ribavirin for an additional 24 hours. Viral supernatants were harvested by freeze-thaw as above and titered by tissue culture infectious dose.

### R-selection through serial passage

For each passage, HeLa cells were plated the day prior to infection in 6 well plates at a density of 7.25 x 10^5^ cells per well, yielding 1.2 x 10^6^ cells on the day of infection. Infections were initiated with passage 3 stocks of either WT or 3D^G64S^, and each passage was performed at an MOI of 0.5 (6 x 10^5^ TCID_50_ units) in 1ml of media for one hour with occasional rocking. After one hour, the inoculum was aspirated, the cells were washed twice with PBS, and 2ml of fresh growth media applied. For the first 15 passages, WT and 3D^G64S^ viruses were harvested at 4 and 4.5 hours, respectively. For passages 16-50, WT and 3D^G64S^ viruses were harvested at 3.5 and 4 hours, respectively. Control populations were infected in the same manner except that viruses were harvested at 24 hours post infection. There were (5) r-selected WT lineages, (5) r-selected 3D^G64S^ lineages, (5) control WT lineages, and (5) control 3D^G64S^ lineages. Viruses were titered at every 5^th^ passage to maintain an MOI of approximately 0.5.

### Selection and identification of second-site suppressors of RdRp variant 3D^K359H^

The 3D^K359R^ RdRp has slower polymerization kinetics and higher fidelity relative to WT [64]. The 3D^K359H^ RdRp has similar characteristics (see Table S4). HeLa cells were transfected by electroporation with viral RNA transcript, added to HeLa cell monolayers and incubated at 37°C. After two days the media were passaged onto a separate monolayer of HeLa cells. Upon cytopathic effect, viruses were harvested by three repeated freeze-thaw cycles, cell debris was removed by centrifugation, and viral supernatants were titrated. In this time frame, the titer increased approximately 40-fold (from 5.1 x 10^5^ pfu/mL to 2.1 x 10^7^). Viral RNA was isolated with QIAamp viral RNA purification kits (Qiagen), according to the manufacturer’s instructions. The 3Dpol cDNA was prepared from purified viral RNA by RT-PCR and sequenced. The I331F and P356S substitutions were identified together in one experiment, and the P356S substitution was identified in a second (see Table S4).

### In vitro assays of RdRp function

All mutations were introduced into the pET26Ub-PV 3D [65] or pSUMO-PV-3D [66] bacterial expression plasmids using either overlap extension PCR or QuickChange Site-Directed Mutagenesis. The presence of the desired mutations and the absence of additional mutations were verified by DNA sequencing. PV 3Dpol RdRps were expressed and purified as described previously [65,66]. The sym-sub assays used to measure assembly/elongation kinetics of purified RdRp on a defined template were performed as described in [6,30,67]. All assays had 1 μM primer/template and 2 μM enzyme.

### Adaptability of WT and 3D^G64S^

For HeLa cells, adaptability was measured using the 24 hour passage control lineages from the r-selection experiment. The fitness values of WT and 3D^G64S^ populations were measured by competition assay, as above, using samples from passages 0, 5, 10, 15, and 20. For PVR 3T3 cells, serial passages were performed as follows. Cells were seeded in 6 well plates at a density of 7.6 x 10^5^ cells per well the day prior to infection, yielding approximately 1 x 10^6^ the day of infection. Serial passage lineages were initiated with passage 1 stocks of either WT or 3D^G64S^, and each passage was performed at an MOI of 0.5 in 1ml for one hour. After one hour, the inoculum was aspirated, the cells were washed twice with PBS, and 2ml of fresh growth media applied. Viruses were harvested at 24 hours and titered every 4^th^ passage to ensure an MOI of 0.5. There were (5) replicate lineages of WT and 3D^G64S^.

### Infection of transgenic mice

This study was approved by the Institutional Animal Care and Use Committee at the University of Michigan and is compliant with all relevant ethical regulations. Six to 9 week old mice were used for all experiments. The age range and distribution of males and females in each group for each experiment are reported in Table S2. On the day of each infection, a general health exam was performed on all animals by University veterinary technical staff and animals were assigned unique ear tag identifiers. For survival analyses, mice were infected intramuscularly with 50μl to each hindlimb for the total dose of 100μl. Mice were observed twice daily for lethargy, hunched posture, scruffy fur, paralysis, or decreased mobility and euthanized when they exhibited bilateral hindlimb paralysis. Over 90% of all assessments were performed by members of the University veterinary technical staff, who were blinded to the hypotheses and expected outcomes of the studies. All surviving animals were euthanized after 12 days. These endpoints were also used to calculate the PD50 using the Spearman-Karber method (Table S3).

For tissue distribution analyses, mice were infected intravenously via tail vein with 100μl and observed twice daily as above. Mice were euthanized for severe morbidity or on day 5, the conclusion of the experiment. Whole organs were isolated from all mice and homogenized in PBS using a Bead Beater. The homogenates were clarified by centrifugation at 15,800 x g for 4 minutes in a microfuge and the supernatant extracted with chloroform. Half of this supernatant was titered by TCID_50_. RNA was extracted from the remainder using Trizol.

### Next generation sequencing

We amplified poliovirus genomes as four overlapping cDNA by RT-PCR. RNA was harvested from either cell free supernatants or tissues as above and was reverse transcribed using the SuperScript III First Strand Synthesis System for RT-PCR (Invitrogen 18080) and a mixture of random hexamers and oligo dT primer. The four genomic fragments were amplified using primer pairs: WFP37 FORWARD BASE 1 5’ TTAAAACAGCTCTGGGGTTGTACC 3’ + WFP41 REVERSE BASE 2434 5’ GCGCACGCTGAAGTCATTACACG 3’; WFP39 FORWARD BASE. 1911 5’ TCGACACCATGATTCCCTTTGACT 3’ + WFP42 REVERSE BASE 4348 5′ AATTTCCTGGTGTTCCTGACTA 3′; WFP13 FORWARD BASE 4087 5′ ATGCGATGTTCTGGAGATACCTTA 3′ + WFP43 REVERSE BASE 5954 5′ CCGCTGCAAACCCGTGTGA 3′; WFP40 FORWARD BASE 5545 5′ TTTACCAACCCACGCTTCACCTG 3′ + WFP33 REVERSE BASE 7441 5′ CTCCGAATTAAAGAAAAATTTACCCC 3′. The thermocycler protocol was: 98°C for 30 sec then 30 cycles of 98°C for 10 sec, 68°C for 20 sec, 72°C for 3 min, followed by a single cycle of 72°C for 5 min, then 4°C hold. For each sample, amplification of all 4 fragments was confirmed by gel electrophoresis and equal quantities of each PCR product were pooled. Seven hundred fifty nanograms of each cDNA mixture were sheared to an average size of 300 to 400bp using a Covaris S220 focused ultrasonicator. Sequencing libraries were prepared using the NEBNext Ultra DNA library prep kit (NEB E7370L), Agencourt AMPure XP beads (Beckman Coulter A63881), and NEBNext multiplex oligonucleotides for Illumina (NEB E7600S). The final concentration of each barcoded library was determined by Quanti PicoGreen dsDNA quantification (ThermoFisher Scientific), and equal nanomolar concentrations were pooled. Residual primer dimers were removed by gel isolation of a 300-500bp band, which was purified using a GeneJet Gel Extraction Kit (ThermoFisher Scientific). Purified library pools were sequenced on an Illumina MiSeq with 2 x 250 nucleotide paired end reads. All raw sequence data have been deposited at the NCBI short read archive (Bioproject PRJNA396051, SRP113717).

### Variant detection

Sequencing reads that passed standard Illumina quality control filters were binned by index and aligned to the reference genome using bowtie [68]. Single nucleotide variants (SNV) were identified and analyzed using DeepSNV [69], which relies on a clonal control to estimate the local error rate within a given sequence context and to identify strand bias in base calling. The clonal control was a library prepared in an identical fashion from the pEW-M plasmid and was sequenced in the same flow cell to control for batch effects. True positive SNV were identified from the raw output tables by applying the following filtering criteria in R: (i) Bonferonni corrected p value <0.01, (ii) average MapQ score on variant reads >20, (iii) average phred score on variant positions >35, (iv) average position of variant call on a read >62 and <188, (v) variant frequency >0.001. We only considered SNV identified in a single RT-PCR reaction and sequencing library for samples with copy number ≥10^5^ genomes/μl supernatant or in two separate RT-PCR reactions and sequencing libraries for samples with copy number 10^3^−10^5^ genomes per μl (for example in tissue studies). Our strategy for variant calling as well as our benchmarked sensitivity and specificity are described in [46] and all code can be found at https://github.com/lauringlab/variant_pipeline.

### Statistical Analysis

No explicit power analyses were used in designing the experiments. In most cases we used 5 biological replicates. In a few cases, we used fewer (3) or more (7) where the variance was either sufficiently low or high. The number of replicates, the statistical tests used, and the relevant p values are reported in each Figure legend or the main text (Fig. 2C and 2D only). All replicates within the dynamic range of each assay are reported (i.e. no replicate experiments were excluded). Data on the relative adaptability of WT and 3D^G64S^ populations were analyzed with a 3-level linear mixed effects model estimating a random slope and intercept of time nested within each fitness measurement replicate (measID), nested within each lineage replicate (repID). Virus was included as a fixed effect. Models were fit with the R package lme4, all code for this model can be found at https://github.com/lauringlab/speed_fidelity.

## Acknowledgements

We thank Ashley Acevedo, Leonid Brodsky, and Raul Andino for providing the raw data on poliovirus mutational fitness effects; Santiago Elena and Rafael Sanjuan for helpful suggestions; Carla Pretto, Rajni Sharma, Jacob Perryman and Alexandre Martinez for technical assistance. This work was supported by R01 AI118886 to ASL and R01 AI 45818 to CEC. RW was supported by K08 AI119182 and JTM was supported by the Michigan Predoctoral Training Program in Genetics (T32 GM007544).

## Supporting Information Model 1 – Speed fidelity trade off

The premise of the speed-fidelity tradeoff is that the outcome of competition between two viral strains will be determined by two opposing forces – the speed with which the genome can be replicated and the error rate of replication. The faster genome replication happens, the more errors that occur and the greater the mutational load. In this scenario an optimal competitive fitness will be achieved exactly where the increase in fitness is counterbalanced by decrease in fitness from excess mutational load. Here we present a simple mathematical model to demonstrate this tradeoff, and fit the model to the experimental data.

We start with the classical estimation by Haldane (1937) of the equilibrium mean population fitness, w, as a function of the genomic deleterious mutation rate (U_d_) in units of deleterious mutations per genome per generation.

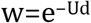

The relationship is shown graphically, here.

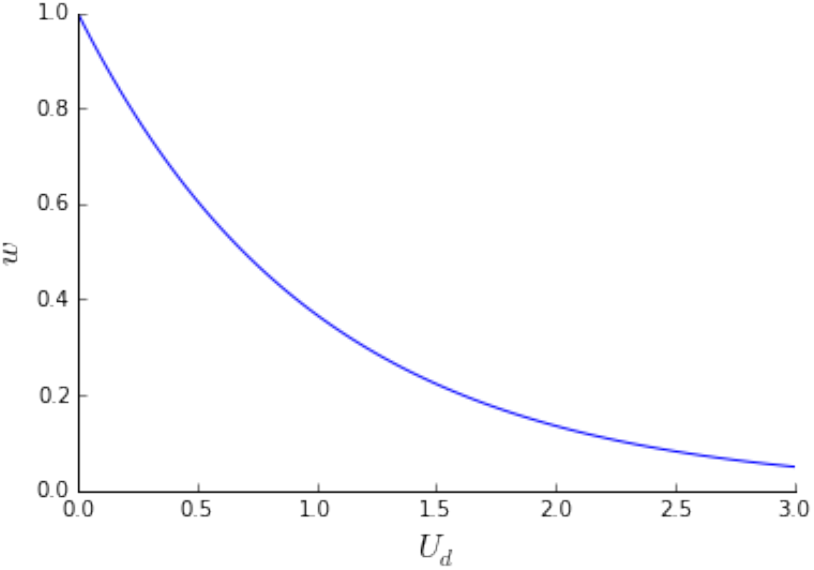

Now we consider competition between two strains that differ in both their speed, c, and their fidelity (as manifest by a deleterious mutation rate, U). If you have two strains, a and b, then the relative fitness of a to b will be:

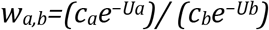

Where c_a_ and c_b_ are the genome replication rates and U_a_ and U_b_ are the deleterious genomic mutation rates per genome per generation for strains a and b, respectively.

To understand the effect of a mutagen on the relative fitness of one strain to another, we add a mutation rate multiplier (mu external, σ), which multiplicatively modifies the baseline mutation rate. We constrain σ to take values greater than one.

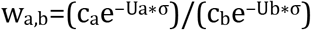

As shown below, as the mutation rate multiplier goes up, the fitness of a high fidelity variant increases relative to wild type (WT). The plot on the left shows both strains (WT and a high fidelity variant together). The equilibrium, where the two strains have equal fitness, is at a sigma value where the two lines cross (as indicated by the arrow, right)

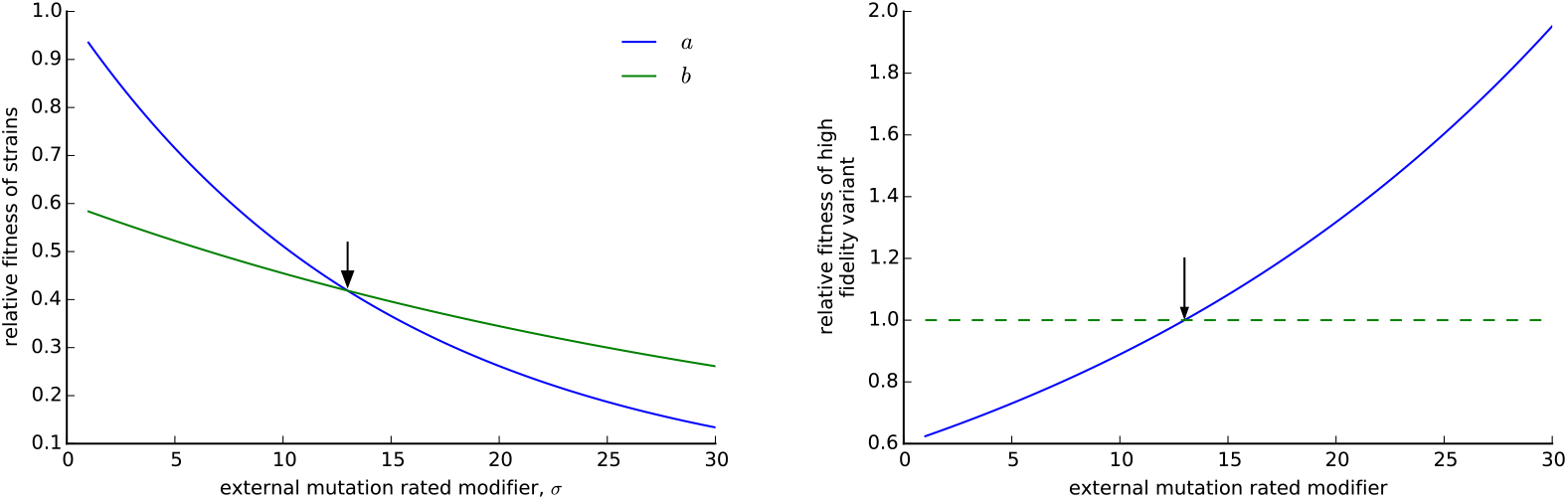

With this simple model, we can use the empirical estimates of mutation rates and relative fitness values over a range of ribavirin concentrations (which increase the mutation rate multiplier, σ) to estimate the deleterious mutation rate and the amount of mutation load experienced by WT poliovirus and the 3D^G64S^ high fidelity variant. The data are presented in Figures 1E and 1F and available in the annotated Jupyter notebook available at https://github.com/lauringlab/speed_fidelity.

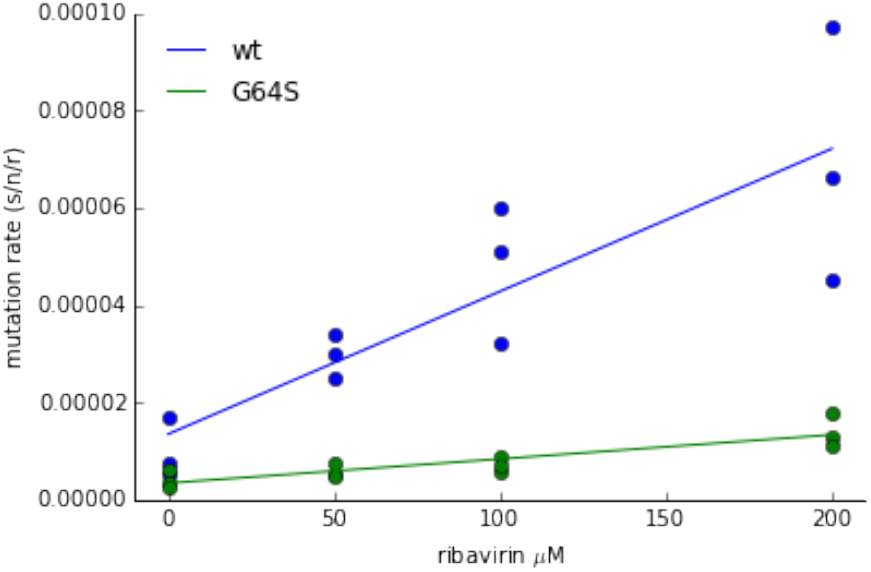

The G64S mutation has two effects (see also Fig. 1F). The mutation results in an increase in fidelity, represented as a downward shift in the line. The mutation also results in relative resistance to ribavirin, which manifests as a decreased slope. Both of these need to be included to estimate the mutational load experienced by WT and the 3D^G64S^ mutant replicating in the presence of increasing mutagen. Linear regression fit to both the WT and 3D^G64S^ mutation rate produce a good fit (r^2^ of .73 and .76 respectively) that is highly statistically significant (p < 0.001 for both). The fit of the linear regression was used to estimate the mutation rate for WT and 3D^G64S^ in the absence of ribavirin (1.34 x 10^−5^ s/n/r and 3.43 x10^−6^ s/n/r respectively) and at the point of equilibrium.

We solved for the two unknown variables: (i) *n*, the number of deleterious sites in the genome, and (ii) *c*, the fitness cost of the G64S mutation in the absence of mutational load, which is relative to the wild type (arbitrarily set to 1). Because there are two unknown variables, we need two equations to solve for them. We use the measured relative fitness in the absence of ribavirin and the ribavirin concentration where the relative fitness is expected to be 1 (150μM, see Fig. 1E). We used the mutation rates estimated for each strain and ribavirin concentration (μstrain,conc). At 0μM ribavirin, the fitness of 3D^G64S^ relative to WT was measured as 0.67, which gives us our first equation:

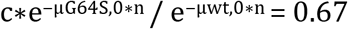

Next, we looked at the competitive fitness data and used the point at which the two strains have equal fitness (approximately 150μM of ribavirin, see Fig. 1E):

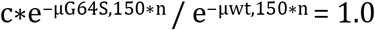

Now, with two equations, we solved for the two unknown values (n and c). Assuming they have equal fitness at 150μM, the fitness cost of 3D^G64S^ absent the cost from mutational load is 0.60. The effective number of sites with deleterious mutations is 10959 (48% of all possible mutations, given 3 possibilities at every site). The fitness cost due to mutation load that is experienced by WT in absence of ribavirin is 0.137. The fitness cost due to mutational load experienced by 3D^G64S^ in absence of ribavirin is 0.037. These relationships are shown graphically below, where the dashed line indicates the fitness in the absence of mutational load, the shaded area indicates the effect of mutational load on fitness and the solid lines indicate the overall effect on fitness.

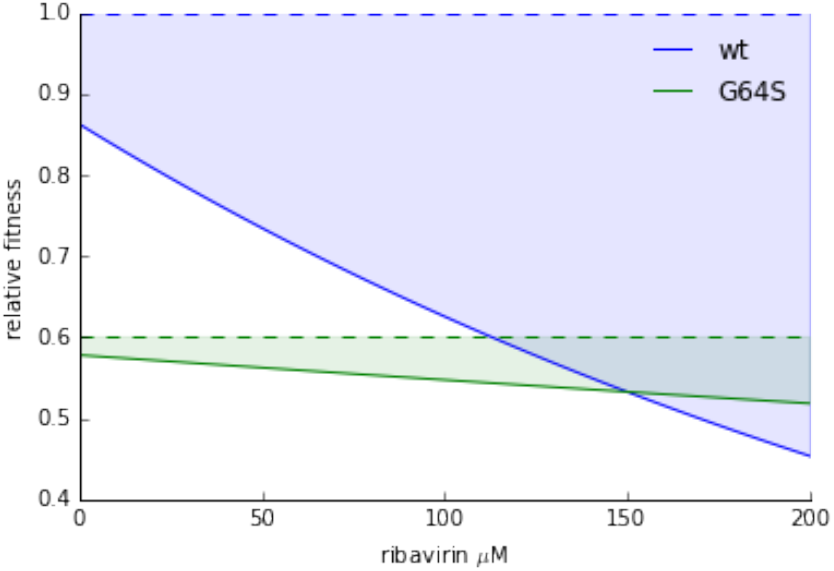

## Supporting Information Model 2 – Within host bottleneck

We developed the following model to measure the effective bottleneck that restricts poliovirus populations between the site of inoculation and the central nervous system. We applied a simple probabilistic model to data described in [48]. In this study, 27 mice were infected with 2 x 10^7^ PFU of poliovirus (2-5 fold higher than the LD_50_ in this particular mouse model). The inocula consisted of 4 subpopulations at equal concentrations, each tagged with a neutral sequence bar code. In separate experiments, the authors showed that all 4 bar codes were present at the site of infection and that all four bar codes were capable of replicating simultaneously in the brain. Rarely were all 4 bar codes present in the brain following infection, suggesting that the populations were subject to within host bottlenecks. Similar results were observed for IV and IP routes of infection. In fact, IM appeared to be the least stringent mode of infection. To estimate the bottleneck between the site of infection and the brain, we modeled the infection process as a random sampling event. This assumption was justified as: (i) there is no evidence that a “jackpot” mutation is needed to enter the central nervous system. (ii) the bar codes were selectively neutral. (iii) all bar codes were equally likely to be present in the brain.

The probability of a sample size of *n* containing *K* unique types given there are *N* total unique types available (all present at equal frequency) derives from discrete probability theory and is given by

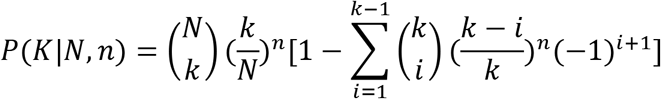

(see for example, Ross SM. 2010. *A First Course in Probability*. Prentice Hall, pages 121-122). In terms of the above experimental design, we are interested in the probability a subset of size *n* containing 1,2,3, or 4 barcodes (*K*) given 4 possible barcodes (*N*).

Maximum likelihood optimization revealed that a bottleneck of 4 PFU best matched the data. However, this model was constrained, in part, by the fact that no smaller bottleneck could account for the presence of 4 barcodes in the CNS even though this rarely occurred. Indeed simulations revealed this model predicted a higher average number of barcodes than experimentally observed. In particular the model underestimated the probability of only 1 barcode infecting the CNS.

To account for experimental variability in bottleneck sizes between mice we allowed the bottleneck to vary according to a zero-truncated Poisson distribution. We truncated the Poisson distribution because at the high doses used in the original experiment and replicated in this current work poliovirus entered the CNS in all infected mice. That is, there were no mice with zero barcodes.

When we allow *n*, the bottleneck size, and in our model, to follow a zero-truncated Poisson distribution parameterized with λ. The likelihood of observing *k* barcodes given *λ* is

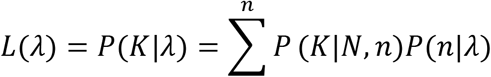

Where *P*(*k*|*N, n*) is our expression above and 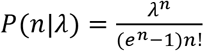 or the probability of n in a zero-truncated Poisson with a parameter *λ*. We approximated the infinite sum above with a partial sum of the first 100 terms as we expected a small bottleneck, and the probability of an *n* of 50 with *λ* = 100 is on the order of 10^−10^ and negligible. We then searched for the *λ* that maximized the sum of the log of this likelihood, which was calculated for each mouse. We found that a *λ* of 2.44 with a 95% confidence interval of (1.39 − 3.82) best fit the data. The mean of a zero-truncated Poisson is given by 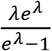. Therefore the mean bottleneck size is 2.67.

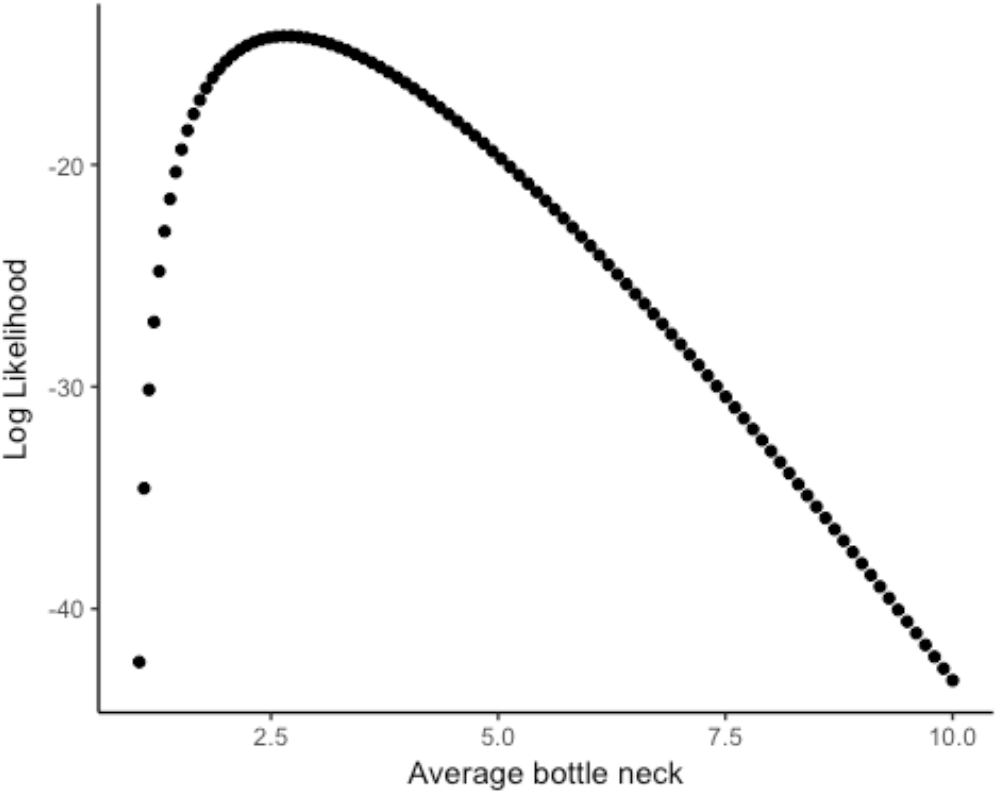

To test the fit we performed 10,000 simulations. Each simulation included 27 mice and each mouse had a bottleneck size drawn from a zero-truncated Poisson with a *λ* of 2.44. For illustration we also simulated the data with an average bottleneck of 10. The output of the simulations is shown below and in Fig. 4A. The shaded regions represent the area occupied by 95% of the simulations with the dark regions representing the interquartile range of the simulations.

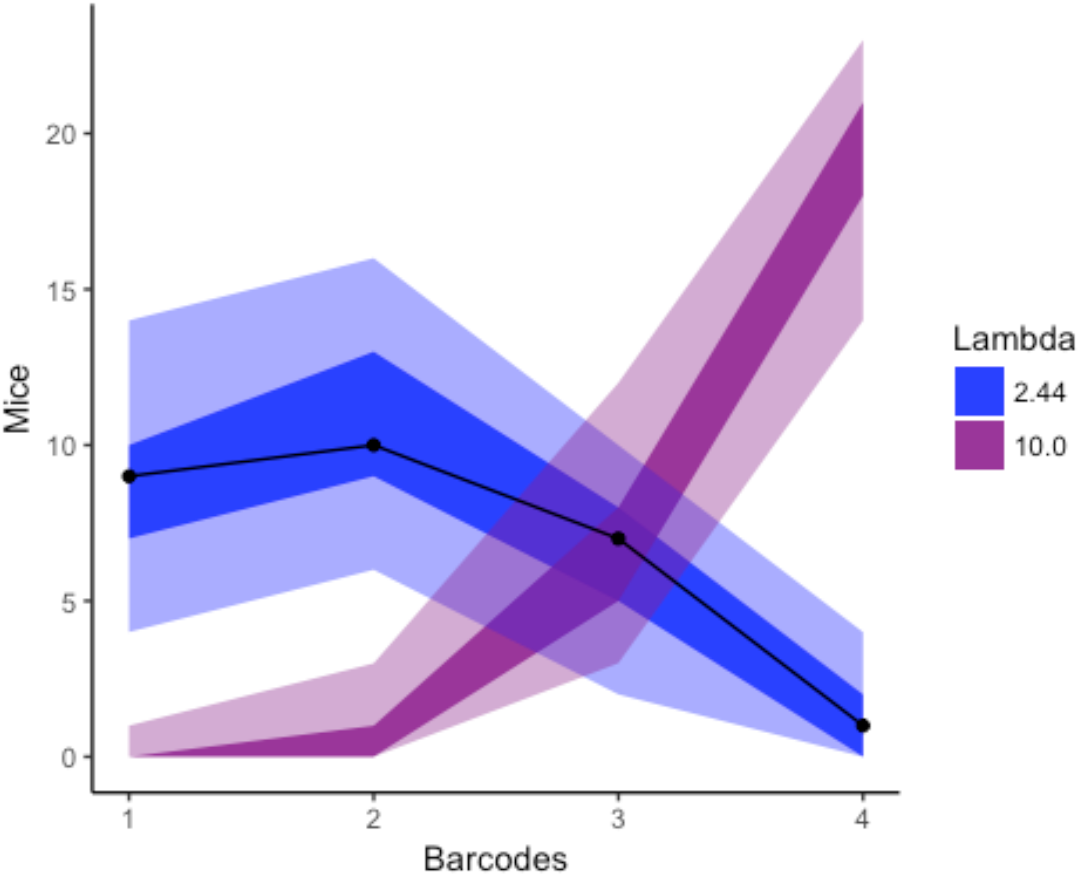

We also checked the fit by asking how the output of the simulations matched the actual experimental data (e.g. how often did we see 9 mice with 1 bar code, 10 with 2, 7 with 3, and 1 with 4).

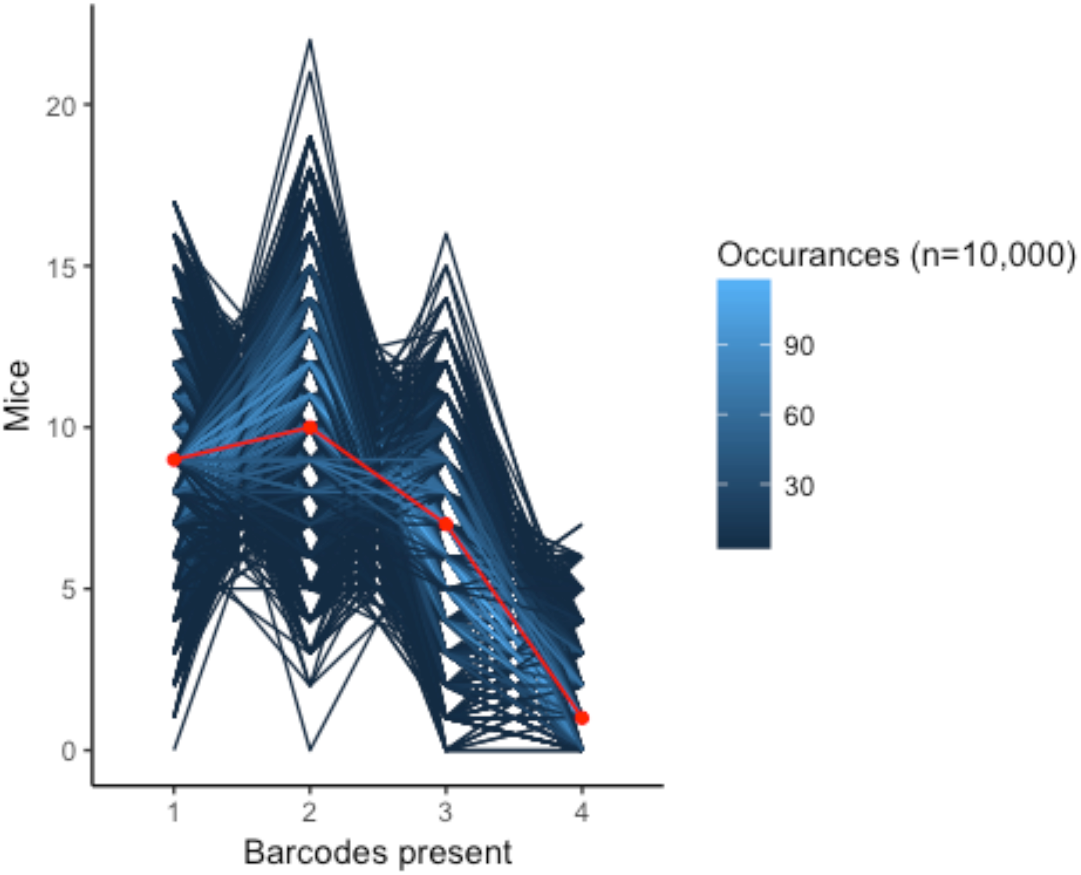

In contrast, we did not replicate the data once in 10,000 simulations with a bottleneck of 10.

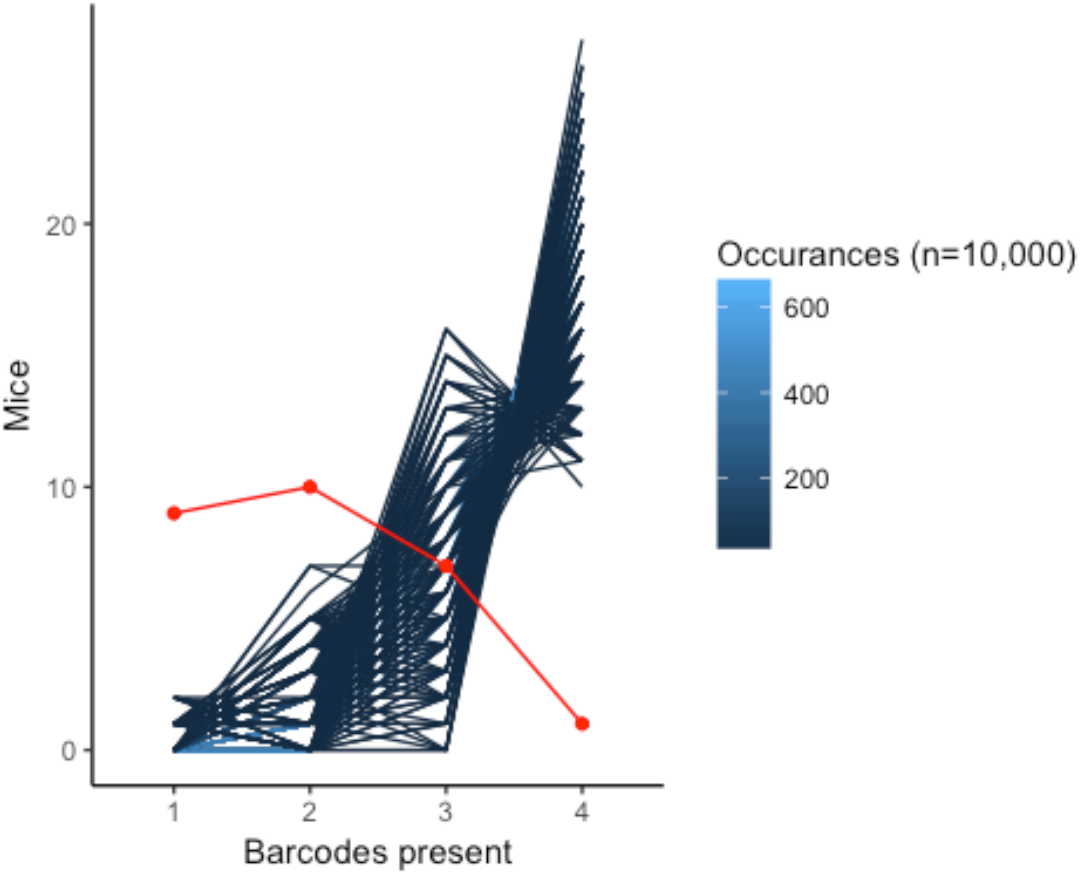

While we modeled the infectious process from inoculation to invasion of the CNS as a single sampling procedure, this process is made up of several bottlenecks imposed on the virus as it passes through different body compartments [49]. We, therefore, interpret our mean bottleneck of 2.67 as an aggregate, within-host bottleneck.

Population bottlenecks have been show to be dose dependent. However, it is likely that the inoculating dose modeled here, and used in the presented work, saturates this dose dependency. For example, Pfeiffer et al. only observed a slight increase in the average number of barcodes present in the CNS when mice were inoculated with 100x more PFU. In fact, the bottleneck appears to only be overcome when very young (two week old) mice were inoculated.

**Table S1.** Mutations conferring resistance to 1mM guanidine. Shown are results from 15 plaques from populations treated with no drug (WT) or 200μM ribavirin (R).

| Plaque | Mutations (position, nucleotide change, amino acid change) | Plaque<br>200µM<br>R | Mutations (position, nucleotide change, amino acid change) |
| --- | --- | --- | --- |
| WT2 | 4614 U->A, F->Y | R1 | 4676 G->U, A->S |
| WT6 | 4614 U->A, F->Y | R8 | 4676 G->U, A->S |
| WT12 | 4614 U->A, F->Y | R10 | 4676 G->U, A->S |
| WT14 | 4676 G->U, A->S | R12 | 4676 G->U, A->S |
| WT1 | 4682 A->U, M->L | R13 | 4676 G->U, A->S |
| WT3 | 4682 A->U, M->L | R7 | 4363 C->U, VAL(SYN) ; 4682 A->C, M->L |
| WT4 | 4682 A->U, M->L | R3 | 4702 G->A, M->I |
| WT5 | 4682 A->U, M->L | R2 | 4802 A->C, I->L |
| WT9 | 4682 A->U, M->L | R4 | 4802 A->C, I->L |
| WT10 | 4682 A->U, M->L | R5 | 4802 A->C, I->L ; 4810 C->U,PRO(SYN) |
| WT11 | 4682 A->U, M->L | R6 | 4797 G->A,SER(SYN) ; 4820 G->A, A->U |
| WT7 | 4702 G->A, M->I | R9 | 4820 G->U, A->S |
| WT13 | 4820 G->U, A->S | R11 | 4459 A->G, I->M ; 4820 G->U, A->S |
| WT8 | 4442 A->G,U->A ; 4823 C->A,H->N | R14 | 4820 G->U, A->S |
| WT15 | NONE in 2C | R15 | 4823 C->A, H->N |

**Table S2.**
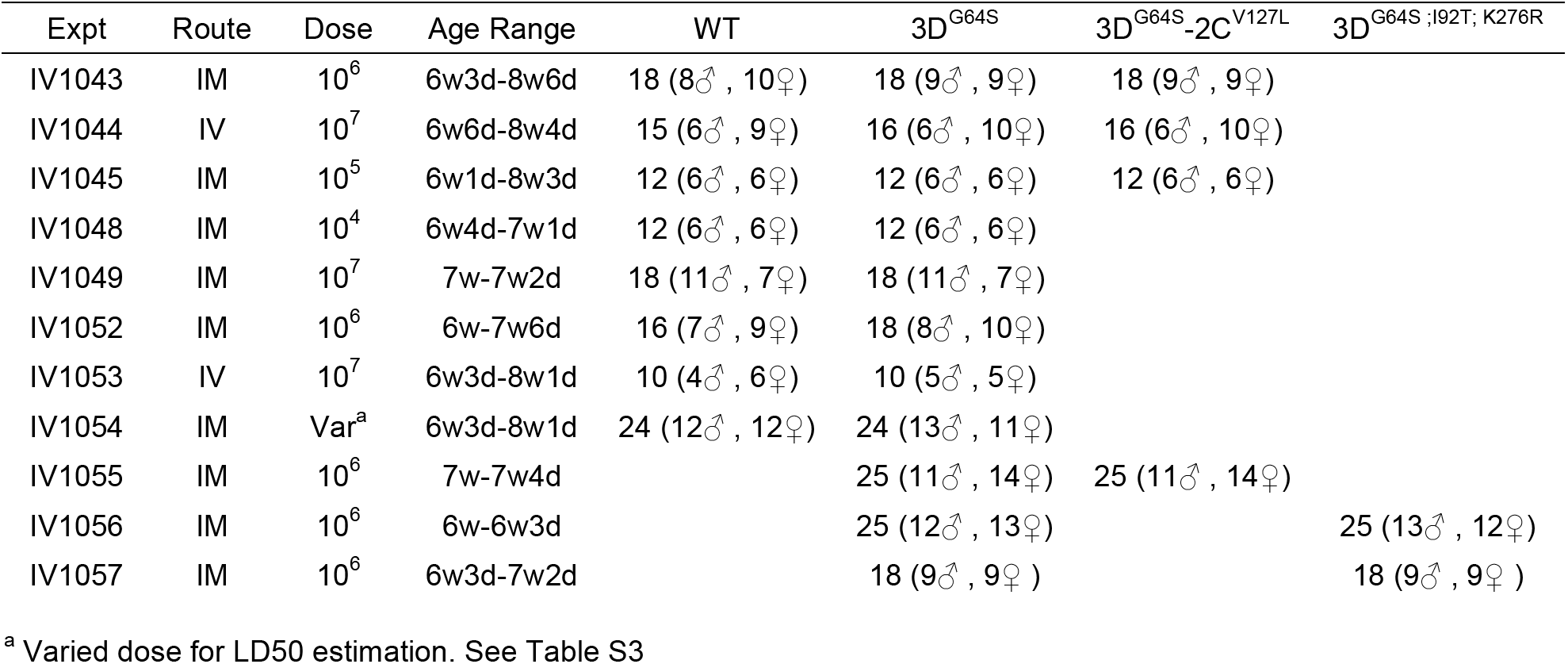
Number, age, and sex of mice used in all experiments.

| Expt | Route | Dose | Age Range | WT | 3D<sup>G64S</sup> | 3D<sup>G64S</sup>-2C<sup>V127L</sup> | 3D<sup>G64S ;I92T; K276R</sup> |
| --- | --- | --- | --- | --- | --- | --- | --- |
| IV1043 | IM | 10<sup>6</sup> | 6w3d-8w6d | 18 (8♂ , 10♀) | 18 (9♂ , 9♀) | 18 (9♂ , 9♀) |  |
| IV1044 | IV | 10<sup>7</sup> | 6w6d-8w4d | 15 (6♂ , 9♀) | 16 (6♂ , 10♀) | 16 (6♂ , 10♀) |  |
| IV1045 | IM | 10<sup>5</sup> | 6w1d-8w3d | 12 (6♂ , 6♀) | 12 (6♂ , 6♀) | 12 (6♂ , 6♀) |  |
| IV1048 | IM | 10<sup>4</sup> | 6w4d-7w1d | 12 (6♂ , 6♀) | 12 (6♂ , 6♀) |  |  |
| IV1049 | IM | 10<sup>7</sup> | 7w-7w2d | 18 (11♂ , 7♀) | 18 (11♂ , 7♀) |  |  |
| IV1052 | IM | 10<sup>6</sup> | 6w-7w6d | 16 (7♂ , 9♀) | 18 (8♂ , 10♀) |  |  |
| IV1053 | IV | 10<sup>7</sup> | 6w3d-8w1d | 10 (4♂ , 6♀) | 10 (5♂ , 5♀) |  |  |
| IV1054 | IM | Var<sup>a</sup> | 6w3d-8w1d | 24 (12♂ , 12♀) | 24 (13♂ , 11♀) |  |  |
| IV1055 | IM | 10<sup>6</sup> | 7w-7w4d |  | 25 (11♂ , 14♀) | 25 (11♂ , 14♀) |  |
| IV1056 | IM | 10<sup>6</sup> | 6w-6w3d |  | 25 (12♂ , 13♀) |  | 25 (13♂ , 12♀) |
| IV1057 | IM | 10<sup>6</sup> | 6w3d-7w2d |  | 18 (9♂ , 9♀ ) |  | 18 (9♂ , 9♀ ) |

<sup>a</sup> Varied dose for LD50 estimation. See Table S3

**Table S3.** Raw data for calculation of LD50 (PD50, as paralysis triggered euthanasia per protocol). Calculation and output were obtained using tsk package in R with no trimming. Similar values were obtained using the logit method.

|  | 10<sup>2</sup> pfu | 10<sup>3</sup> pfu | 10<sup>4</sup> pfu | 10<sup>5</sup> pfu | 10<sup>6</sup> pfu | 10<sup>7</sup> pfu | PD50 | 95% CI |  |
| --- | --- | --- | --- | --- | --- | --- | --- | --- | --- |
| WT | 0/6 | 0/6 | 3/6 | 5/6 | 6/6 | 6/6 | 1.47E+05 | 4.7E+04 | 4.6E+05 |
| 3D<sup>G64S</sup> | ND | 0/6 | 1/6 | 3/6 | 6/6 | 6/6 | 6.81E+05 | 2.2E+05 | 2.1E+06 |

Age range 6w3d-8w1d; 25♂ , 23♀
All animals dosed intramuscularly
PD50 by Spearman-Karber method

**Table S4.** Kinetic parameters for nucleotide incorporation and misincorporation for purified RdRp. The k_pol_ for the correct nucleotide measures the speed of polymerization in vitro. The k_pol,corr_/k_pol,incorr_ is an in vitro surrogate for fidelity, as it measures the relative rates of incorporation for the correct and incorrect nucleotides. A higher ratio indicates higher fidelity.

| RdRp | Correct Nucleotide |  | Incorrect Nucleotide |  |
| --- | --- | --- | --- | --- |
| | $k_{pol} \text{ (s}^{-1}\text{)}$ | $K_{d,app} \text{ (}\mu\text{M)}$ | $k_{pol} \text{ (s}^{-1}\text{)} [\times 10^{-3}]^a$ | $k_{pol,corr.} / k_{pol,incorr.}$ |
| WT | $37 \pm 3$ | $180 \pm 40$ | $4.2 \pm 0.6$ | 8,800 |
| K359H | $4.0 \pm 0.2$ | $250 \pm 20$ | $0.10 \pm 0.02$ | 40,000 |
| I331F K359H | $8.5 \pm 0.1$ | $170 \pm 10$ | $0.74 \pm 0.13$ | 11,000 |
| P356S K359H | $9.8 \pm 0.3$ | $110 \pm 10$ | $0.78 \pm 0.09$ | 13,000 |
| I331F P356S K359H | $17 \pm 1$ | $120 \pm 10$ | $4.9 \pm 0.8$ | 3,500 |

<sup>a</sup> Determined at saturating concentrations of nucleotide substrate.

**Figure S1.**
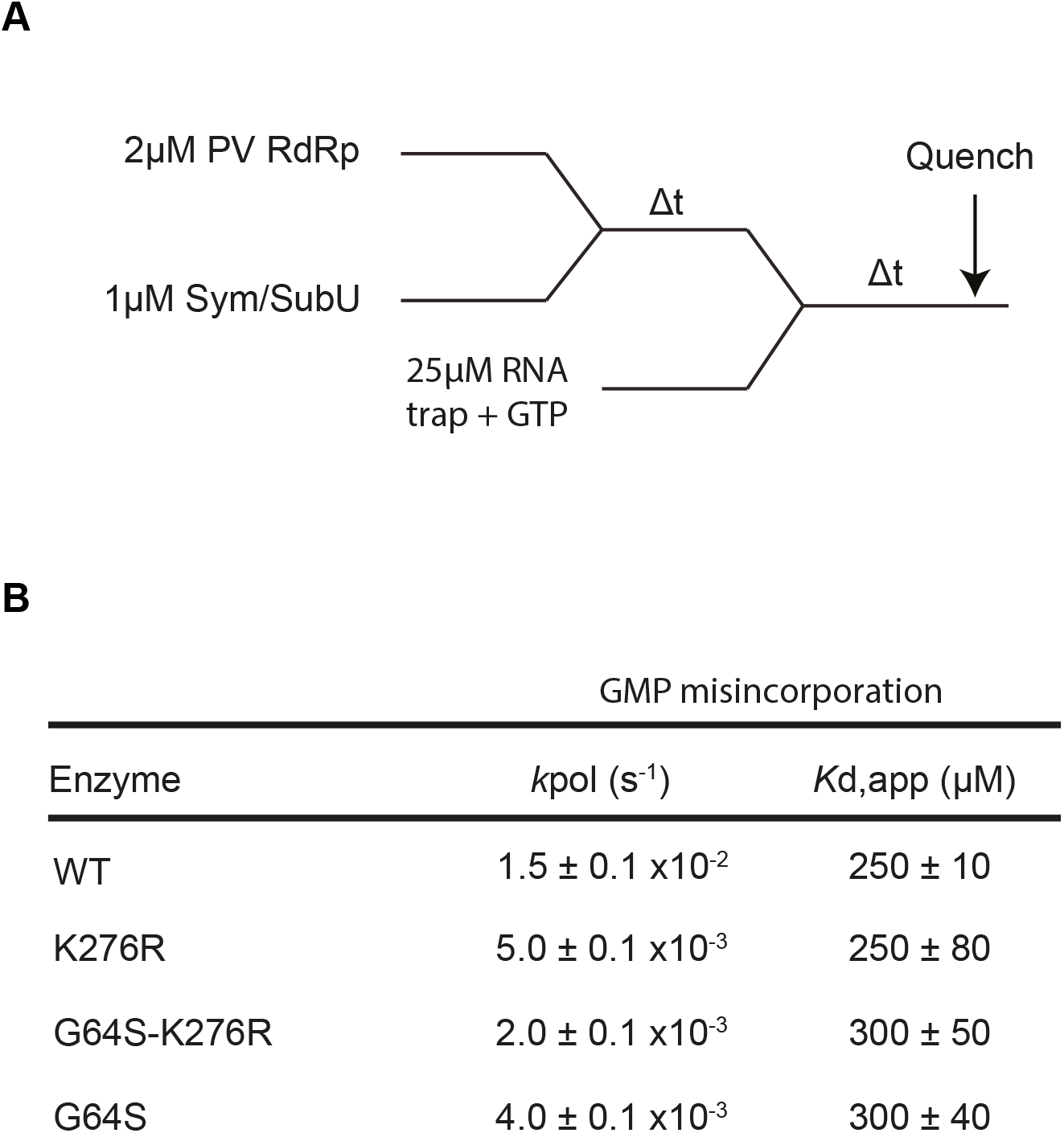
In vitro assay of polymerase mediated single nucleotide incorporation. (A) Schematic of GTP misincorporation assay (G opposite the U). Primer-template (sym-subU) and polymerase are assembled in the absence of nucleotide. GTP and a 25-fold excess of unlabeled trap RNA are then added after an incubation period (Δt). Excess of RNA trap ensures that if the polymerase dissociates from primer-template it is taken up by the trap and cannot re-assemble. (B) Kpol and Kd for GMP misincorporation. Concentrations of GTP were used over time-points to calculate Kpol and Kd. Each GTP time-course was plotted to single exponential, then combined plot to hyperbola. Note that all polymerase variants have the I92T mutation.

**Figure S2.**
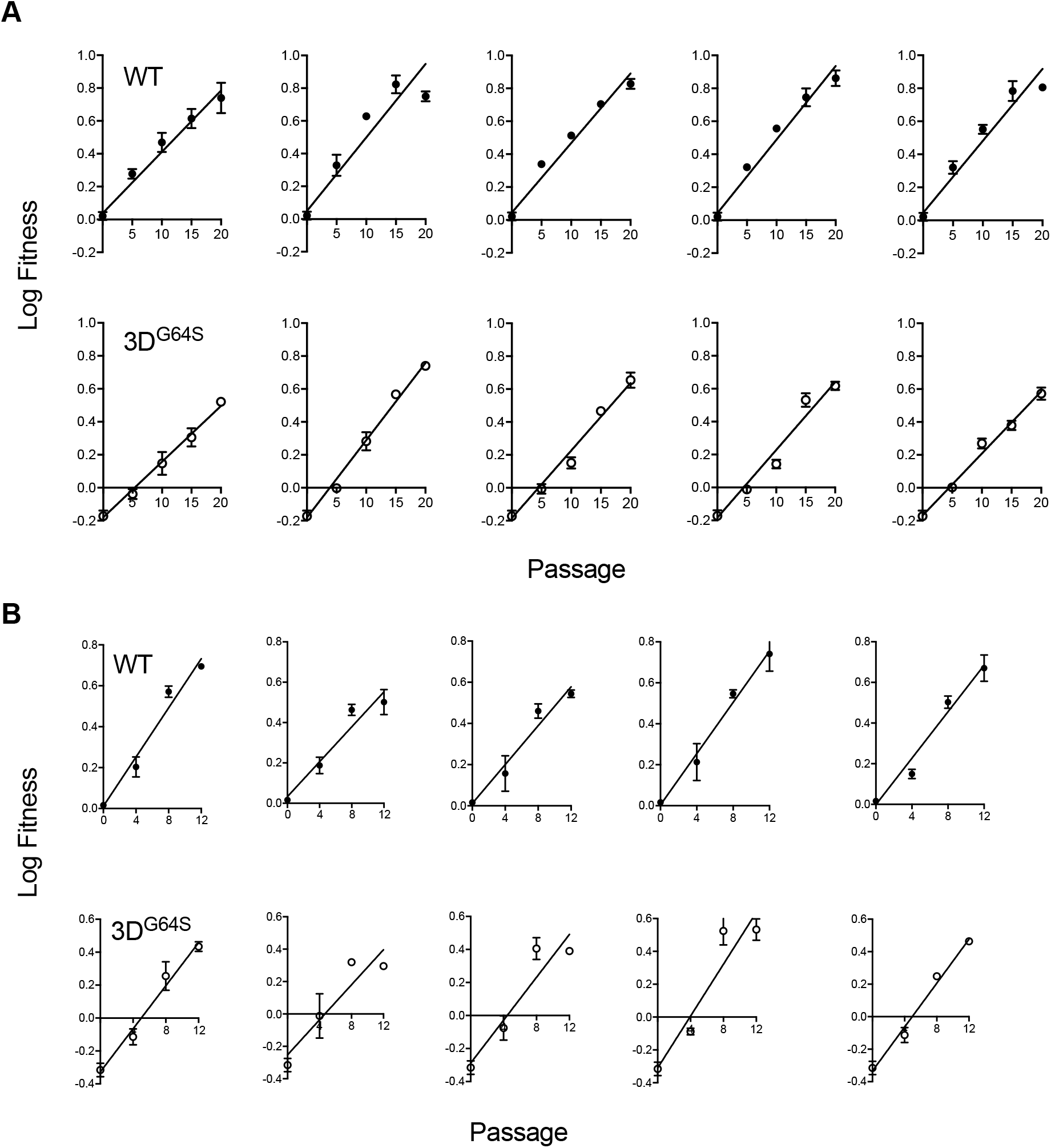
Log fitness versus passage in adaptability experiment. Each point is the fitness mean ± standard deviation of three replicate competition assays. (A) WT and 3DG64S on HeLa (B) WT and 3DG64S on PVR-3T3. For the graphs in (B), the relationship of log fitness vs. passage was not linear past passage 16, and only passages 1-12 are shown.

**Figure S3.**
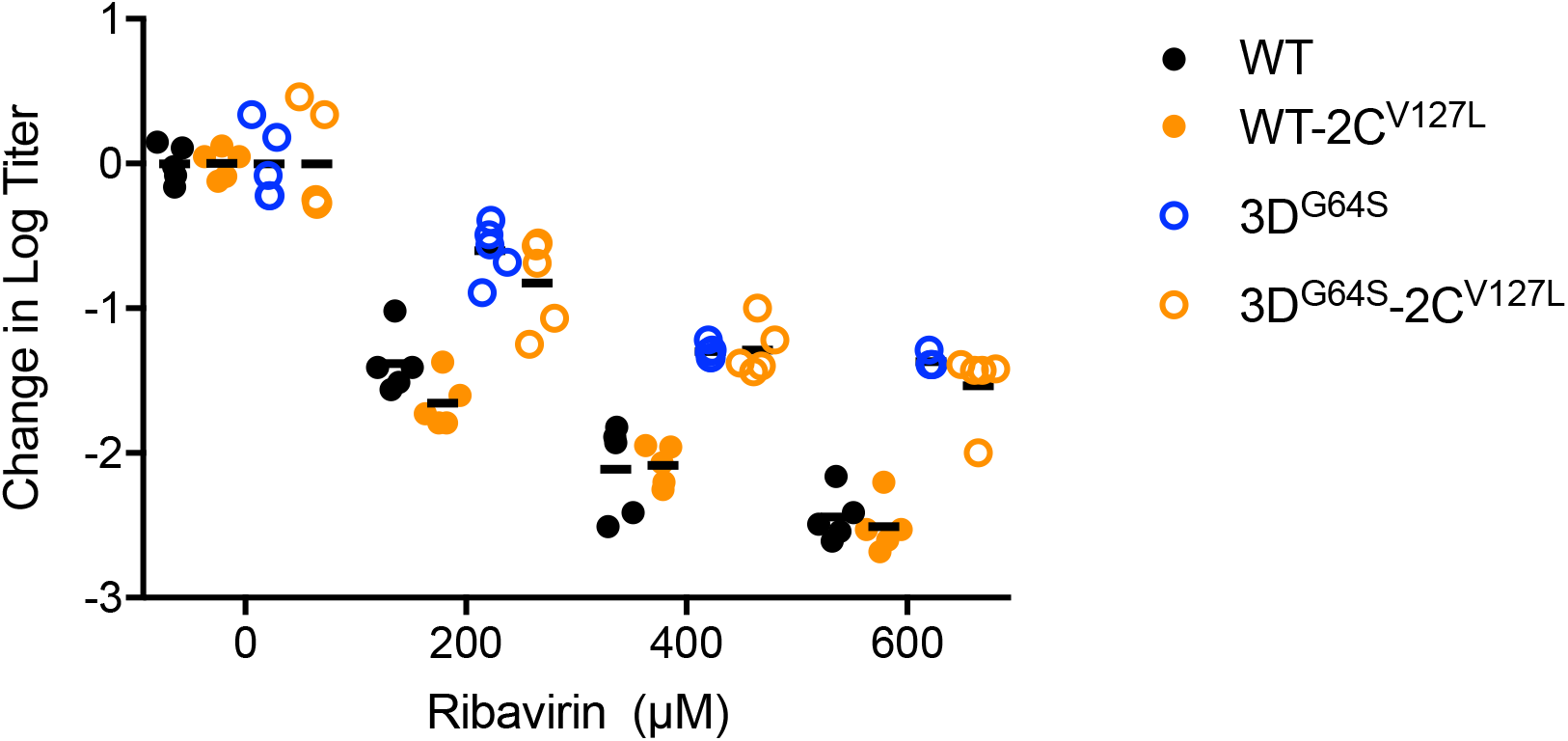
Mutagen sensitivity of 2C-V127L variants. HeLa were infected at an MOI of 0.1 with the indicated viruses in presence of various concentrations of ribavirin. After 24 hours, titers of mock and ribavirin-treated populations were determined by TCID50 and those of ribavirin-treated populations were normalized to mock-treated controls (mean of 5 measurements per virus). Shown are the changes in titer (y-axis, mean, 5 replicates) for each virus at each drug concentration (x-axis). A greater reduction in titer (more negative number) indicates higher mutagen sensitivity and suggests a higher baseline mutation rate.

